# Cancer therapy with a CRISPR-assistant telomerase-activating gene expression system

**DOI:** 10.1101/291302

**Authors:** Wei Dai, Xinhui Xu, Danyang Wang, Jian Wu, Jinke Wang

## Abstract

Telomerase is silent in most normal somatic cells while active in 90% of tumor cells. Various telomerase activity inhibitors have been developed to treat cancer but all failed due to side effects. Here we acted oppositely to develop a cancer therapy named telomerase-activating gene expression (Tage) by utilizing the telomerase activity in tumor cells. By using CRISPR/Cas9 functions, the Tage system can effectively kill various cancer cells, including HepG2, HeLa, PANC-1, MDA-MB-453, A549, HT-29, SKOV-3, Hepa1-6, and RAW264.7, without effecting normal cells. By using homothallic switching endonuclease and adeno-associated virus, the Tage system realizes its *in vivo* application. The virus-loaded Tage system can significantly and specifically kill the cancer cells in mice by intravenous drug administration without side effects or toxicity.

**One Sentence Summary:** Killing cancer cells in body with a gene therapy missile detonated by telomerase.

## Main Text

The human telomere consists of many repetitive TTAGGG sequences, which functions in chromosome protection, positioning and replication (*1*, *2*). Telomeres terminate with a 3′ single-stranded overhang that is bound by multiple variant proteins, which is essential for both telomere maintenance and capping (*3*). The eukaryotic DNA replication machinery can’t replicate the extreme end of chromosomes, causing gradually shortened chromosomes along with cell division. In order to repair the damaged telomere in some special cells, a specialized protein named as telomerase has been developed. Telomerase is a natural RNA-containing enzyme that can synthesize the repetitive telomere sequences (*4*), which thus helps to maintain the integrity of the genome in some cells such as embryonic stem cells (*5*). Telomerase is silenced in most normal tissue cells but reactivated in most human cancer cells (*6*, *7*), suggesting that telomerase may be a good target to cancer therapy (*8*, *9*). Therefore, many telomerase inhibitors has been developed to treat cancers; however, none of them becomes the applicable clinical drugs due to their side effects (*9*-*11*).

In this study, we act oppositely to utilize the telomerase activity in cancer cells to develop a cancer therapy named as the telomerase-activated gene expression (Tage) (Fig.1). In this system, an effector gene expression vector (effector for short) that carries a telomerase-recognizable 3ʹ single-stranded sequence (5ʹ-TTAGGG-3ʹ) is transfected into cancer cells so that it can be elongated by telomerase, which will produce a synthesized double-stranded telomere repeat sequence at the end of effector. Simultaneously, a vector able to express an artificial transcription factor (TF), dCas9-VP64/sgRNA, is cotransfected in cancer cells, which can produce the dCas9-VP64/sgRNA complex that can recognize and bind the telomerase-synthesized double-stranded telomere repeat sequences at the end of effector. This binding can thus activate the expression of an effector gene, Cas9, on effector. The expressed Cas9 protein can associate with the telomere-targeting sgRNA (TsgRNA) produced by the dCas9-VP64/sgRNA vector. The Cas9/TsgRNA complex then damages chromosomes of cancer cells by cutting their telomeres, which will induce the apoptosis or death of cancer cells. However, in normal tissue cells that lack the telomerase activity, the Tage system can’t be activated and thus exerts no effect on cells.

**Fig.1.**
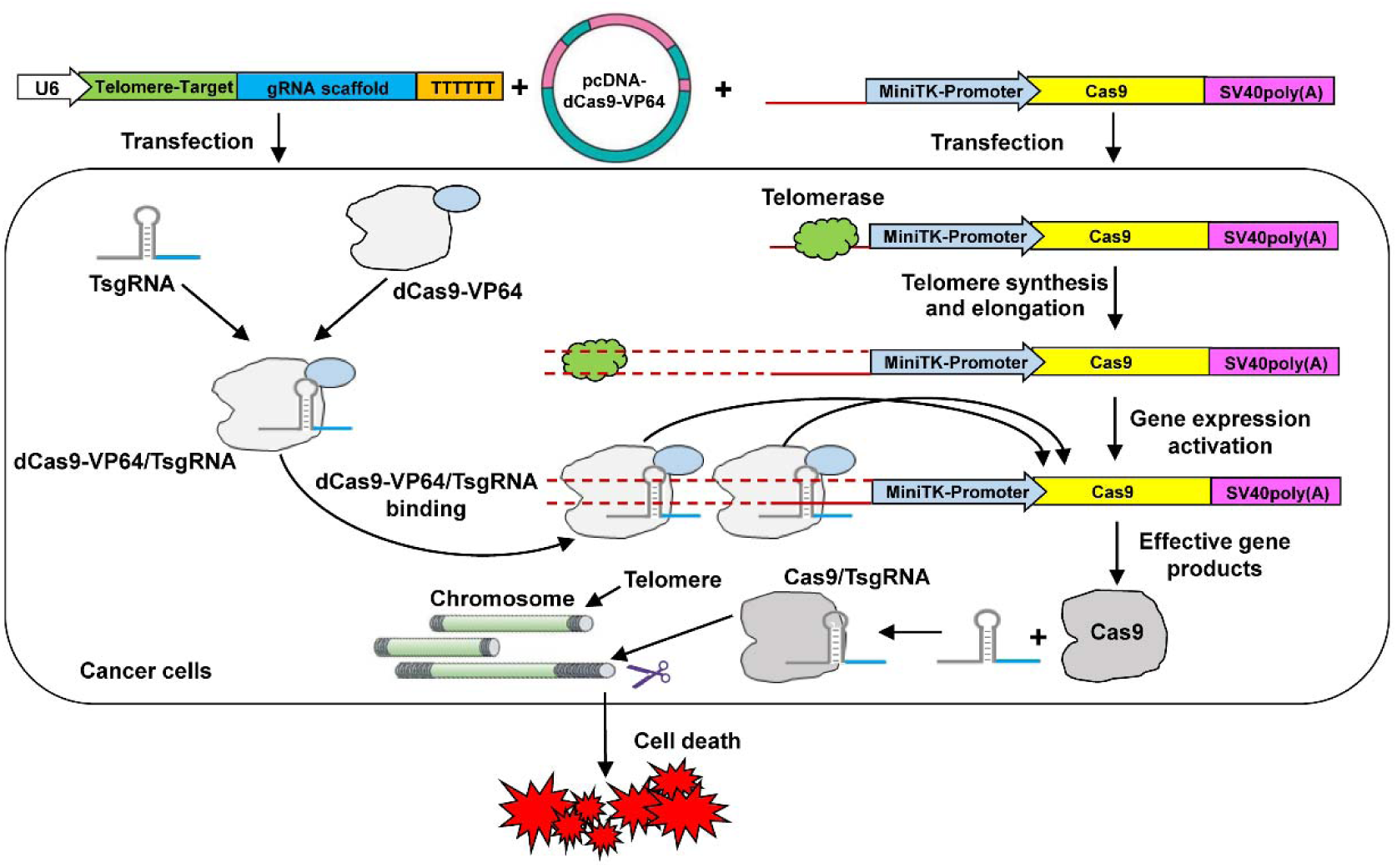
Schematic of Tage system to kill tumor cells.

The synthesis of telomere repetitive sequence at the end of effector is a key step for the Tage system coming into its effect. To verify the process, we transfected cells with the Tage system and extracted the genomic DNA (gDNA) of cells after cultivated for 24 h. We detected the gDNA by polymerase chain reaction (PCR) using the TS and CX primers of telomere repeat amplification protocol (TRAP)(*12*). The results reveal that the gDNA of telomerase-positive cells (293T and HepG2) can be amplified; however, the gDNA of telomerase-negative cells (MRC-5 and HL7702) can’t be amplified (Fig.S1, lane 1 and 2). Additionally, the gDNAs of cells transfected with an effector with blunt telomeric sequence also can’t be amplified (Fig.S1, lane 3). These results indicate that the single-stranded telomeric sequence at the end of effector can only be elongated by telomerase in cancer cells.

We then verify the effectiveness of the Tage system with a reporter construct. We transfected cells with a Tage system including an effector in which the ZsGreen was used as effector gene (sTMEP). The results indicate that the ZsGreen was successfully expressed by the 293T and HepG2 cells (Fig.2A and 2B), but not by the MRC-5 and HL7702 cells (Fig.2C and 2D). Moreover, in all cells transfected by various DNAs as controls, the ZsGreen protein was not expressed (Fig.S2A-2D). Especially, even the 293T and HepG2 cells were transfected with an blunt-end effector (bTMEP), the ZsGreen protein still can’t be expressed (Fig.S2A and 2B). These results reveal that the Tage system can only be activated by telomerase in cancer cells. Without the single-stranded telomerase recognizable telomeric sequence at the end of effector, the telomeric DNA can’t be synthesized. These results also suggest that the newly synthesized telomeric DNA at the end of effector was double-stranded as previously testified (*13*); otherwise, the effector gene also can’t be expressed in cancer cells because dCas9/sgRNA only bind double-stranded DNA.

We next used Cas9 as effector gene of Tage system to aim at killing cancer cells by cutting telomere. We conceived that the TsgRNA can guide the Cas9 protein to the telomeres of chromosomes, where Cas9/TsgRNA can cut telomeric DNA and induce cell apoptosis. We thus transfected cells with an effector expressing Cas9 (sTMCP) together with vectors expressing TsgRNA and dCas9-VP64. The results reveal that the Tage system can induce the significant death of all transfected cancer cells, including HepG2, HeLa, PANC-1, MDA-MB-453, A549, HT-29, SKOV-3, Hepa1-6, and RAW264.7 (Fig.3). However, the Tage system exerts no effects on normal cells MRC-5 and HL7702 (Fig.3). Moreover, even the cancer cells were transfected by an effector with a blunt end (bTMCP), their viability was not affected (Fig.S3). The effectiveness of Tage system with Cas9 as effector gene was also confirmed by other negative transfections in cancer cells (Fig.S3).

**Fig.2.**
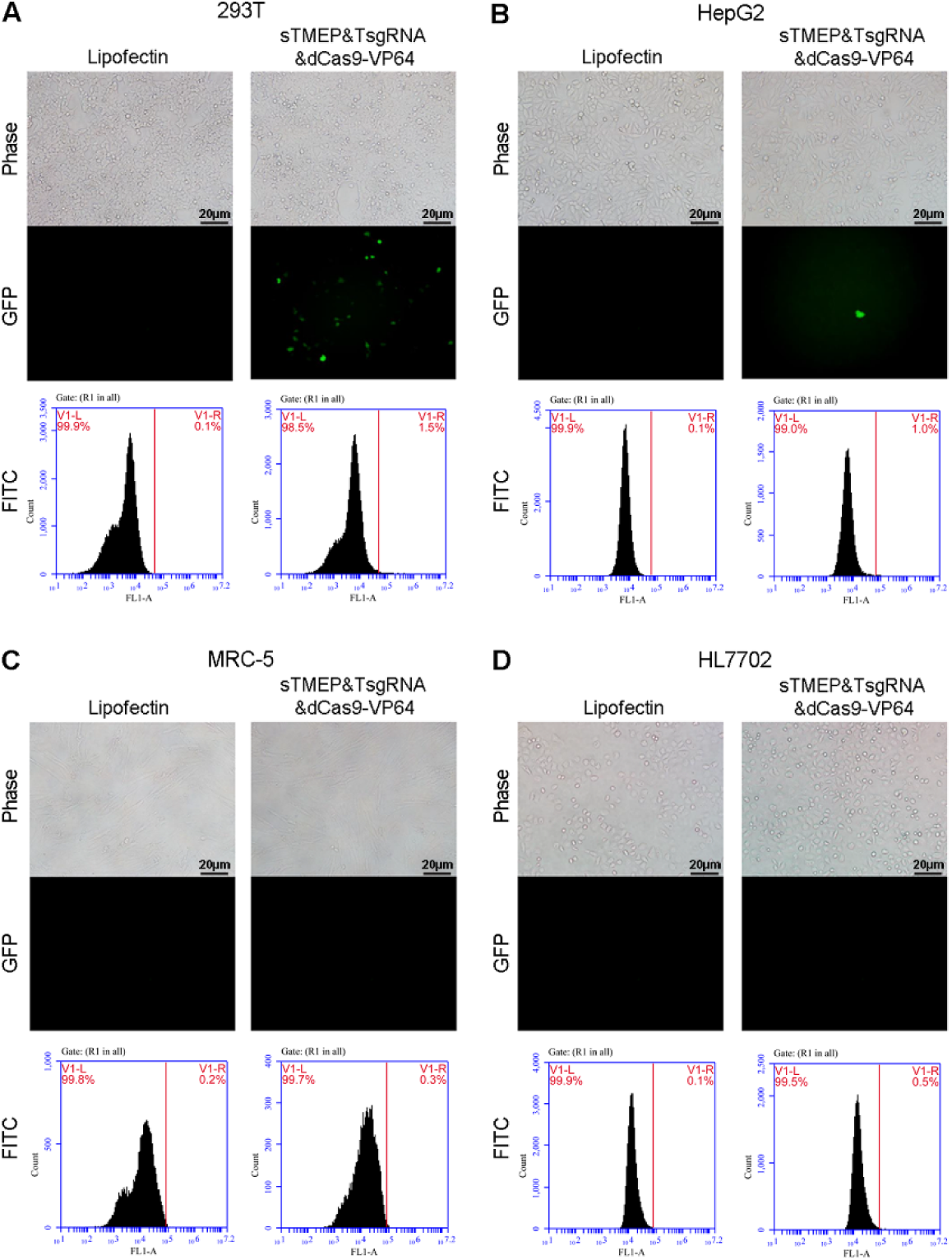
Evaluation the specificity of Tage system in different cell lines. The fluorescence microscope pictures and representative flow cytometry analysis of 293T (A), HepG2 (B), MRC-5 (C), and HL7702 (D) cells, which were transfected by a Tage system with an effector gene ZsGreen.

**Fig.3.**
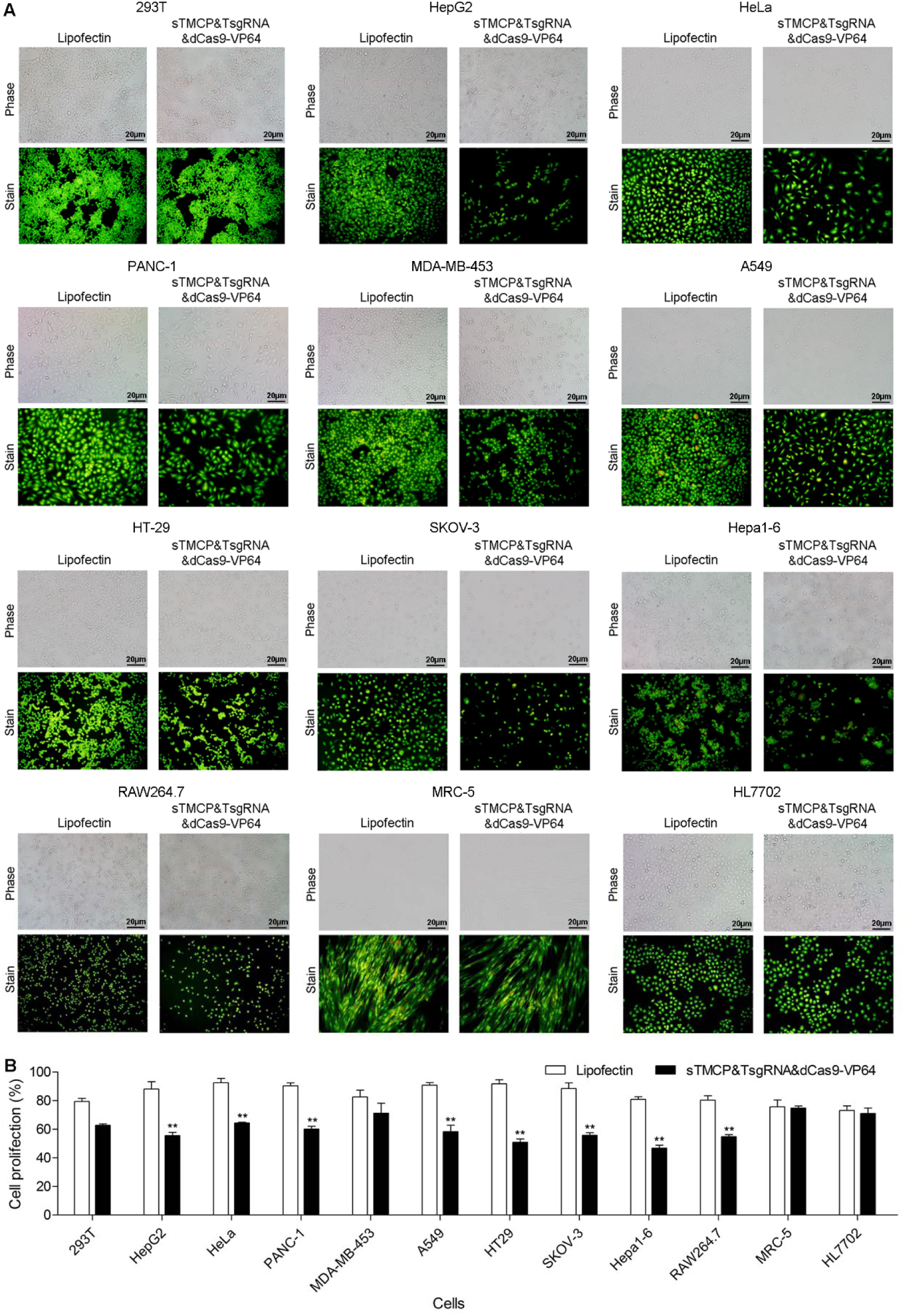
Kill tumor cells with Tage system. (A) The microscope pictures and representative acridine orange staining pictures of various cell lines transfected by the Tage system. (B) The alamar blue assay of cell viability of various cell lines transfected by the Tage system. TsgRNA, telomere-targeting sgRNA.

Although the linear effector with a telomerase recognizable stick end can be easily transfected into cells *in vitro* by lipofectin, it can’t be transfected into *in vivo* cells using virus vectors. We therefore next focus on addressing the issue. We found that it was reported that a four-base overhang, 3ʹ-TTGT-5ʹ, produced by Homothallic switching endonuclease (HO) cutting its target DNA sequence can also be recognized and elongated in *S. cerevisiae* cells by telomerase (*13*). To evaluate whether this 4-bp overhang can also be utilized in human cells by the Tage system, we constructed an effector with a stick end of 3ʹ-TTGT-5ʹ that can express reporter gene ZsGreen (HOsite-sTMEP). We cotransfected cells with HOsite-sTMEP, TsgRNA, and dCas9-VP64. The results indicate that the ZsGreen was expressed by 293T and HepG2 cells (Fig.S4A and 4B), but not by MRC-5 and HL7702 cells (Fig.S4C and 4D). Similarly, the negative transfection controls, especially an effector with a blunt end of HO site (HOsite-bTMEP), did not activate the ZsGreen expression in all transfected cells (Fig.S4). These results indicate that the four-base overhang produced by HO can be used by the Tage system. Therefore, we next constructed another vector that can express HO protein (C1-HO) to explore whether the expressed HO protein can cut a blunt-ended effector containing a HO site (HOsite-bTMEP) to produce a stick end of 3ʹ-TTGT-5ʹ. We thus transfected cells with a new Tage system consists of four vector including C1-HO, HOsite-bTMEP, TsgRNA, and dCas9-VP64. The results reveal that the ZsGreen protein was expressed by the 293T and HepG2 cells (Fig.S4A and 4B), but not by the MRC-5 and HL7702 cells (Fig.S4C and 4D).

To simplify the Tage system of above four vectors, we then constructed a TsgRNA and dCas9-VP64 coexpression plasmid (TsgRNA-dCas9-VP64). We confirmed the plasmid by cotransfecting cells with TsgRNA-dCas9-VP64 and sTMCP. The results indicate that the two-vector Tage system can induce the significant death of cancer cell HepG2 (Fig.S5B), while exerts no effect on normal cells MRC-5 and HL7702 (Fig.S5C and S5D). With the verified TsgRNA-dCas9-VP64 plasmid, we next transfected cells with a Tage system consisting of three vectors, C1-HO, T-HOsite-TMCP, and TsgRNA-dCas9-VP64. The results reveal that the cancer cell HepG2 can be significantly killed by the Tage system (Fig.S6B); however, the growth of normal cells MRC-5 and HL7702 was not affected by the Tage system (Fig.S6C and 6D). We noticed that the growth of cancer cell HepG2 was also not affected by the transfection of a Tage system lacking HO expression vector. These results suggest that the HO expression vector is necessary to the Tage system, which expressed HO protein can cut the T-HOsite-TMCP plasmid to produce a telomerase recognizable stick end. With the three verified vectors including TsgRNA-dCas9-VP64, T-HOsite-TMCP, and C1-HO, it is possible to apply the Tage system to *in vivo* by using the virus vector. It should be noted that the cells transfected with C1-HO together with T-HOsite-TMCP or TsgRNA-dCas9-VP64 did not show death (Fig.S6C). This is very important because the HO expression is driven by stronger promoter CMV. This suggests that the HO enzyme did not damage the gDNA of transfected cells because there is no HO recognizable sites in human genome.

To further verify that the cell death induced by the Tage system is genuinely resulted from the Cas9/TsgRNA-induced DNA damage, we constructed a plasmid TsgRNA-Cas9 that can directly express TsgRNA driven by U6 promoter and Cas9 and EGFP driven by CMV promoter. We transfected cells with the vector and found that it can express EGFP and induce death in all transfected cells, including 293T, HepG2, MRC-5, and HL7702 (Fig.S7).

Finally, we explore the *in vivo* application of the Tage system. For this purpose, we firstly cloned the C1-HO, T-HOsite-TMCP, and TsgRNA-dCas9-VP64 into the adeno-associated virus (AAV). We testified the packaged recombinant AAVs (rAAVs) by cotransfecting cells with the rAAV-HOsite-TMCP, rAAV-TsgRNA-dCas9-VP64, and rAAV-HO (rAAV-TSD). The results reveal that the rAAV-TSD induced significant death of cancer cells HepG2 and Hepa1-6 (Fig.S8, A and B), but exert no effects on normal cells MRC-5 and HL7702 (Fig.S8, C and D). Other control transfections including a blank virus (rAAV-MCS) also did not affect the growth of all transfected cells (Fig.S8).

With the testified rAAV-TSD, we performed animal experiments. In the first animal experiment, the mouse Hepa1-6 cells were firstly mixed with rAAV-TSD and rAAV-MCS *in vitro*, respectively. The cells were then subcutaneously transplanted to two body sides of mice. After two weeks, the tumors with hyperemia grew on both sides of mice transplanted with the Hepa1-6 cells mixed by rAAV-MCS (Fig.4A, Fig.S9A), in the mean size of 29 mm^2^ (Fig.4B). However, the tumors on mice treated by rAAV-TSD were significantly inhibited (Fig.4A, Fig.S9B), in the mean size of 7 mm^2^ (Fig.4B). Most tumors were eradicated in the mice treated by rAAV-TSD (Fig.4A, Fig.S9B).

**Fig.4.**
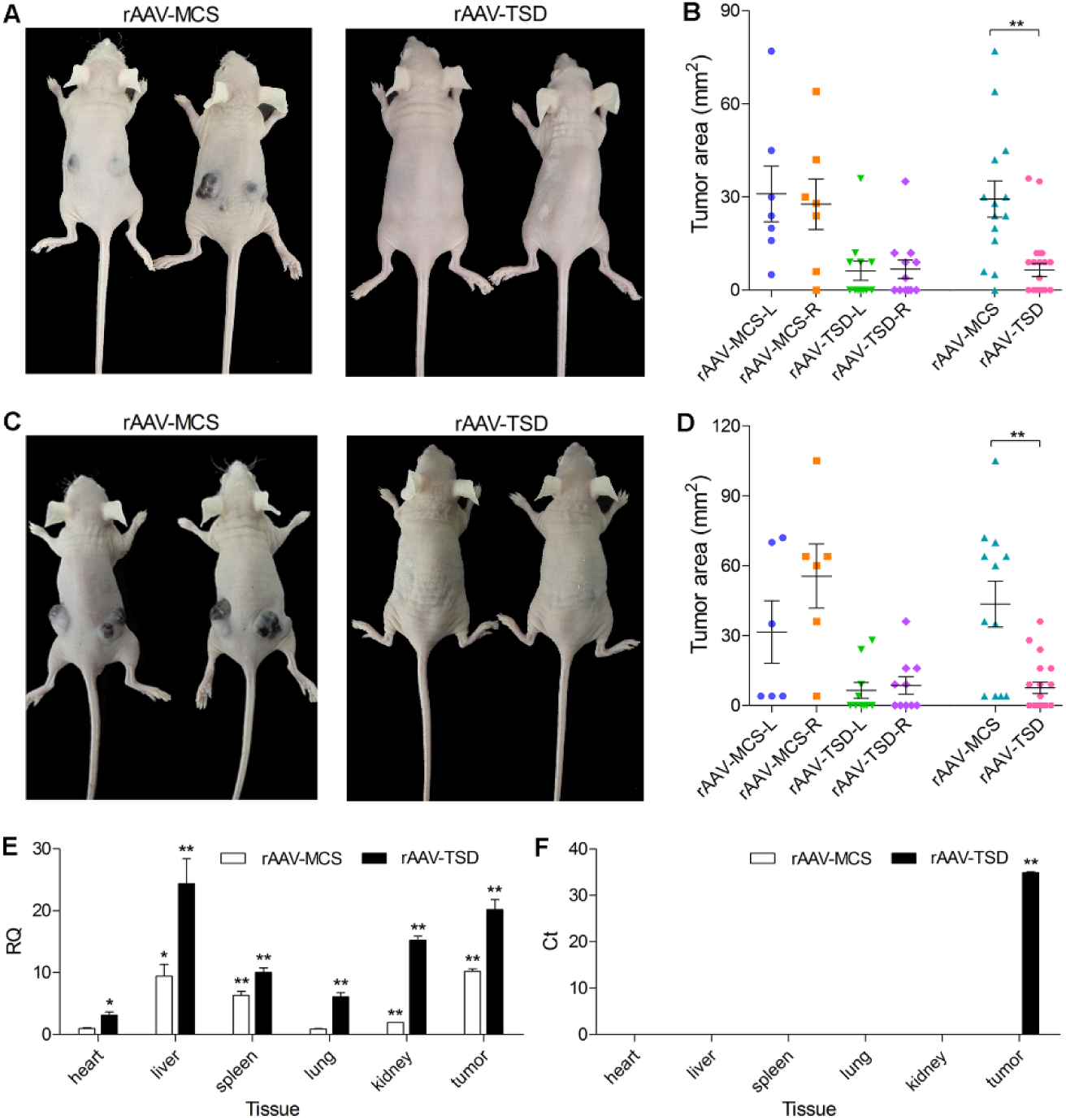
Application of Tage system for cancer therapy *in vivo*. (A) Mice subcutaneously transplanted with Hepa1-6 cells mixed with rAAV-TSD and rAAV-MCS, respectively. (B) Tumor size of mice subcutaneously transplanted with the Hepa1-6 cells mixed with rAAV-TSD and rAAV-MCS, respectively. (C) Tumor-burdened mice intravenously injected with rAAV-TSD and rAAV-MCS, respectively. (D) Tumor size of tumor-burdened mice intravenously injected with rAAV-TSD and rAAV-MCS, respectively. (E) Viral DNA presence in various tissues of tumor-burdened mice intravenously injected with rAAV-TSD and rAAV-MCS, respectively. (F) Cas9 mRNA presence in various tissues of tumor-burdened nude mice intravenously injected with rAAV-TSD and rAAV-MCS, respectively. rAAV-TSD refers to a equivalent mixture of three rAAVs including rAAV-HOsite-TMCP, rAAV-TsgRNA-dCas9-VP64, and rAAV-HO. L, left side; R, right side. \**p ≤ 0.05*, \*\**p ≤ 0.01*.

In the second animal experiment, all mice were firstly subcutaneously transplanted with the Hepa1-6 cells on both sides. After one week, the mice were divided into two groups and intravenously injected with rAAV-MCS and rAAV-TSD, respectively. After one week, the mice were sacrificed for photographing and measuring tumor size. The results revealed that the mice treated with rAAV-MCS grew large tumors on both sides (Fig.4C, Fig.S10A), in the mean size of 44 mm^2^ (Fig.4D). However, the mice treated with rAAV-TSD grew much smaller tumors on both sides (Fig.4C, Fig.S10B), in the mean size of 8 mm^2^ (Fig.4D). Most tumors were eradicated (Fig.4C, Fig.S10B). Importantly, no mice died in two animal experiments after rAAVs were administered, indicating the safety of the used rAVVs.

Finally, to further explore the cancer cell-specific expression of effector gene of the Tage system, we collected the main organs including heart, liver, spleen, lung, kidney, and tumor of the mice of the second animal experiment. We detected the virus DNA and Cas9 mRNA in these tissues with qPCR. The results indicate that the virus presents in all detected tissues with variant abundance (Fig.4E); however, the Cas9 mRNA only presents in tumors (Fig.4F). These results indicate that the Tage system can only be activated in cancer cells, which leads to cancer cell-specific expression of effector gene.

With above *in vitro* cell tests and *in vivo* animal tests, we develop a novel gene therapy of cancers, which depends on a telomerase-activating gene expression technique. This technique fully developed the functions of CRISPR, including the double-stranded cleavage of target DNA (*14*-*16*) and gene expression regulation (*17*, *18*) in cells. The Tage system provides the first *in vivo* application of CRISPR technique in cancer gene therapy without visible side effects or toxicity, which can promote the medical application of gene therapy when it comes of age (*19*).

Clearly, the Tage system is a versatile technical platform that is adaptable. In our view, the telomerase-activating gene expression vector is a powerful missile, whereas the effector gene is its loaded warhead. The warhead is changeable, including “nuclear warhead” like Cas9 used in this study and other “conventional warhead” like variant functional proteins or microRNAs that can lead to cancer cell apoptosis, suicide, reprograming, and differentiation; for example, using the Tage system to express some membrane or secretory proteins intriguing immunotherapy to cancers (*20*). Besides effector gene, the artificial TF in the Tage system can also be changed into other proteins with similar function. For example, using the effector sTMEP, we demonstrate that the artificial TF dCas9-VP64/sgRNA used in the Tage system can be replaced by another artificial TF, transcription activator-like effector (TALE), that targets to the telomeric DNA (Fig. S11).

We used AAV as gene vector in animal experiments. In the past 2017, the US Food and Drug Administration (FDA) has approved several gene therapy such as hemophilia (*21*) and spinal muscular atrophy (*22*). All of these gene therapy clinical tests use AAV as gene vector due to its long-standing safety (*23*). However, a shortcoming of AAV vector is its limited packaging capacity (~5kb). Therefore, we have to package the Tage system into three rAAVs, which can limit the applicable virus dosage and cancer killing effect because the Tage system functions only when three cotransfected rAAVs jointly enter into a cancer cell. However, nanoparticle carriers can be used to slim the Tage system.

## Acknowledgments

This work was supported by the National Natural Science Foundation of China (61571119). Authors declare no competing interests. A patent application has been filed relating to this work.

## Materials and Methods

### Plasmid construction

The plasmid pEGFP-C1 (Clontech) was employed to construct MCS-SV40 polyadenylation signal fragments (Msps) with primers Msps-F and Msps-R, then subcloned into T-clone vector pMD19-T (TaKaRa) to generate plasmid T-Msps. Artificially synthesized telomere and TK miniPRO fragments (TP-F and TP-R) were annealed, extended, and fused with ZsGreen fragments amplified from plasmid pLVX-shRNA-ZsGreen (Addgene) with primers Tm E-F, Tm E-R, ZsGreen-F, and ZsGreen-R, then cloned into plasmid T-Msps using the cutting sites of EcoRI and BamHI enzymes (ThermoFisher Scientific) to generate plasmid T-TMEP. Double-stranded blunt-ended TMEP (bTMEP) fragments were amplified from plasmid T-TMEP with primers Ts-F and Msps-R, and then digested by Nt.BbvCI enzymes (NEB) to generate 3ʹ single-stranded stick-ended TMEP (sTMEP) fragments with primers Ts-R. Two complementary oligonucleotides (TsgRNA-F and TsgRNA-R) with a 20-bp target telomere sequence were annealed then cloned into plasmid px458 (Addgene) using a BbsI digest (NEB) to generate plasmid TsgRNA-Cas9.

Artificially synthesized telomere and TK miniPRO fragments (TP-F and TP-R) were cloned into plasmid T-Msps using the cutting sites of BglII and HindIII enzymes (ThermoFisher Scientific) to generate plasmid T-TMP with primers Tm BglII-F and Tm HindIII-R. Linear Cas9 fragments were amplified from plasmid pCW-Cas9 (Addgene) with primers Cas9 HindIII-F and Cas9 KpnI-R, and then cloned into plasmid T-TMP using the cutting sites of HindIII and KpnI enzymes (ThermoFisher Scientific) to generate plasmid T-TMCP. Double-stranded blunt-ended TMCP (bTMCP) fragments were amplified from plasmid T-TMCP with primers Ts-F and Msps-R, and then digested by Nt.BbvCI enzymes (NEB) to generate 3ʹ single-stranded stick-ended TMCP (sTMCP) fragments with primers Ts-R.

Linear TsgRNA fragments target telomere were amplified from plasmid TsgRNA-Cas9 with primers U6-F and U6-R, then cloned into plasmid pcDNA-dCas9-VP64 (Addgene) using the cutting sites of BglII and XhoI enzymes (ThermoFisher Scientific) to generate plasmid TsgRNA-dCas9-VP64 with primers dCas9 BglII-F, dCas9 SpeI-R, TsgRNA BglII-F and TsgRNA Xhol-R. Two complementary oligonucleotides with homothallic switching endonuclease (HO) sites (Ada HO-F and Ada HO-R) were annealed to generate double-strand DNA (dsDNA) with 4-base overhang (5ʹ-TGTT-3ʹ) on 3ʹ end, then ligated to the BsmBI-predigested TMEP and TMCP fragments for generating the HOsite-sTMEP and HOsite-sTMCP fragments with primers TMEP BsmBI-F and Msps-R, and subcloned into T-clone vector pMD19-T to generate plasmid T-HOsite-sTMEP and T-HOsite-sTMCP respectively. Double-stranded blunt-ended TMEP with HO sites (HOsite-bTMEP) fragments were amplified from plasmid T-HOsite-sTMEP with primers HOsite-TMEP-F and Msps-R. The plasmid C1-HO was constructed by replacing the EGFP gene with the HO genes using the cutting sites of NheI and KpnI enzymes (ThermoFisher Scientific) with primers HO NheI-F and HO KpnI-R.

A linear TsgRNA-dCas9-VP64 fragment were amplified from plasmid TsgRNA-dCas9-VP64 with primers A TsgRNA-F, A TsgRNA-R, A dCas9-F and A dCas9-R, then cloned into plasmid pAAV-MCS vector (Stratagene) using the cutting sites of SalI and BglII enzymes (ThermoFisher Scientific) to generate plasmid pAAV-TsgRNA-dCas9-VP64. Linear HOsite-TMCP fragments were amplified from plasmid T-TMCP with primers A HO-TMCP-F and A HO-TMCP-R, then cloned into plasmid pAAV-MCS vector (Stratagene) using the cutting sites of EcoRI and SalI enzymes (ThermoFisher Scientific) to generate plasmid pAAV-HOsite-TMCP. Linear HO fragments were amplified from plasmid C1-HO with primers A HO-F and A HO-R, then cloned into plasmid pAAV-MCS vector (Stratagene) using the cutting sites of SalI and BglII to generate plasmid pAAV-HO. See Table S1-S5 for full primer sequences and oligos sequences information.

A vector transcription activator-like effector (TALE) targeting telomere DNA sequence (5ʹ-TAGGGTTAGGGTTAGGGT-3ʹ) was constructed with the Golden Gate TALEN and TAL Effector Kit 2.0 (Addgene) according to manufacturer’s instruction.

### Cell culture and transfection

The RAW264.7, Hepa1-6, HEK-293T, HepG2, HeLa, PANC-1, and MDA-MB-453 cells were cultured in Dulbecco’s Modified Eagle Medium (DMEM) (HyClone), supplemented with 10% fetal calf serum (FBS) (HyClone), 100 U/mL penicillin and 100 µg/mL streptomycin. The A549, HT-29, SKOV-3, MRC-5, and HL7702 cells were cultured in Roswell Park Memorial Institute (RPMI) 1640 Medium (HyClone), supplemented with 10% FBS, 100 U/mL penicillin and 100 µg/mL streptomycin. All cell lines were obtained from the cell resource center of Shanghai Institutes for Biological Sciences, Chinese Academy of Sciences and maintained in a humidified incubator set at 37 °C and 5% CO_2_.

DNA was aliquoted into individual tubes before transfection. Cells were seeded into 24-well plates at a density of 0.5×10^5^ cells/well and transfected with various DNA (see following experiments) using Lipofectamine 2000 (ThermoFisher Scientific) using the following protocol. For each transfection, cells were cultured with 500 μL of Opti-MEM (ThermoFisher Scientific) at 37 °C for 30 min when grown to a density of 2×10^5^ cells/well. A stock solution of 50 μL of Opti-MEM, 500 ng of total DNA and 50 μL of Opti-MEM, 2 μL of Lipofectamine 2000 (ThermoFisher Scientific) per transfection was made respectively. The solution was then vortexed respectively and incubated for 5 minutes at room temperature. Thereafter, the Opti-MEM/Lipofectamine solution was added to the individual aliquots of DNA stocked in 50 μL of Opti-MEM, vortexed, and incubated for 20 minutes at room temperature before being added to each well. After incubated for 5 h, the medium of each well was replaced with 500 μL of fresh DMEM or RPMI 1640 medium containing 10% FBS. The cells were further incubated at 37 °C and 5% CO2 for another 24 h. All of these transfections were performed in triplicate.

### Detection of telomerase-synthesized telomeric DNA with PCR

The 293T, HepG2, MRC-5, and HL7702 cells were transfected with sTMEP&TsgRNA&dCas9-VP64, sTMEP, bTMEP, and lipofectin. After 24 h, the genomic DNA (gDNA) was extracted from cells using the TIANamp Genomic DNA Kit (TIANGEN) according to the manufacturer’s instructions. The gDNA was then amplified with PCR using primers TS and CX (Table S1). The PCR reaction was performed in a total volume of 20 µL, containing 10 µL of 2×Premix Taq (TaKaRa), 0.25 µM of primers, and 50 ng of gDNA. PCR profile contained an initial denaturation step at 94 °C for 5 min, followed by 35 cycles of denaturation at 94 °C for 30 s, annealing at 58 °C for 20 s, and elongation at 72 °C for 1 min, then followed by an elongation at 72 °C for 8 min. All PCR amplifications were performed with a blank control at the same time. The PCR products were analyzed in 1.5% agarose gel electrophoresis.

### Test the Tage system using ZsGreen as effector gene

The 293T, HepG2, MRC-5, and HL7702 cells were transfected with various combination of vectors including sTMEP, bTMEP, TsgRNA, dCas9-VP64, and C1-EGFP. After transfection, cells were cultivated for 24 h. Cells were then observed and photographed with a fluorescence microscope (Olympus) at a constant magnification of 200×. The fluorescence intensity of cells were quantified with a flow cytometry (Calibur, BD, USA) and the mean fluorescent intensity (MFI) was calculated by BD software of flow cytometry.

### Test the Tage system using Cas9 as effector gene: killing cancer cells in vitro

The RAW264.7, Hepa1-6, HEK-293T, HepG2, HeLa, PANC-1, MDA-MB-453, A549, HT-29, SKOV-3, MRC-5, and HL7702 cells were transfected with various combination of vectors including sTMCP, bTMCP, TsgRNA, dCas9-VP64 and C1-EGFP. After transfection, cells were cultivated for 24 h. Cells were stained with Acridine orange. Cells were washed twice with PBS, then stained with 100 μg/mL acridine orange (Solarbio) for 10 min at room temperature. Cells were then observed and photographed with a fluorescent microscope (Olympus) at the constant magnification of 200×. For detecting the cell viability with alamar blue, cell lines were seeded into 96-well plates at a density of 1×10^4^ cells/well and cultivated overnight. Cell were then transfected with various combinations of vectors including sTMCP, bTMCP, TsgRNA, dCas9-VP64 and C1-EGFP. After transfection, cells were incubated with 100 μL of fresh medium for 5 h. Cells were then added with 10 μL of alamar blue (YEASEN) per well. Cells were cultivated 24 h and the fluorescence of each well was measured with an excitation wavelength at 530 nm and an emission wavelength at 590 nm using BioTek plate reader (BioTek). The percentage of cell survival in the treated wells was calculated relative to the control-treated wells.

### Test the Tage system containing HO

In the first experiment, the 293T, HepG2, MRC-5, and HL7702 cells were transfected with various combinations of vectors including HOsite-sTMEP, HOsite-bTMEP, TsgRNA, dCas9-VP64, and C1-HO. After transfection, cells were cultivated for 24 h. Cells were then observed and photographed with a fluorescence microscope (Olympus) at a constant magnification of 200×. The fluorescence intensity of cells were quantified with a flow cytometry (Calibur, BD, USA) and the mean fluorescent intensity (MFI) was calculated by BD software of flow cytometry.

In the second experiment, the 293T, HepG2, MRC-5, and HL7702 cells were transfected with various combinations of vectors including sTMCP, TsgRNA-dCas9-VP64, and C1-EGFP. After transfection, cells were cultivated for 24 h. Cells were stained with Acridine orange. Cells were washed twice with PBS, then stained with 100 μg/mL acridine orange (Solarbio) for 10 min at room temperature. Cells were then observed and photographed with a fluorescent microscope (Olympus) at the constant magnification of 200×.

In the third experiment, the 293T, HepG2, MRC-5, and HL7702 cells were transfected with various combinations of vectors including T-HOsite-TMCP, TsgRNA-dCas9-VP64, C1-HO, and C1-EGFP. After transfection, cells were cultivated for 24 h. Cells were stained with Acridine orange. Cells were washed twice with PBS, then stained with 100 μg/mL acridine orange (Solarbio) for 10 min at room temperature. Cells were then observed and photographed with a fluorescent microscope (Olympus) at the constant magnification of 200×.

### Killing cells with Cas9/TsgRNA

The 293T, HepG2, MRC-5, and HL7702 cells were transfected with vectors TsgRNA-Cas9 and C1-EGFP. After transfection, cells were cultivated for 24 h. Cells were stained with Acridine orange. Cells were washed twice with PBS, then stained with 100 μg/mL acridine orange (Solarbio) for 10 min at room temperature. Cells were then observed and photographed with a fluorescent microscope (Olympus) at the constant magnification of 200×. The fluorescence intensity of cells were quantified with a flow cytometry (Calibur, BD, USA) and the mean fluorescent intensity (MFI) was calculated by BD software of flow cytometry. For detecting the cell viability with alamar blue, cell lines were seeded into 96-well plates at a density of 1×10^4^ cells/well and cultivated overnight. Cell were then transfected with vectors TsgRNA-Cas9 and C1-EGFP. After transfection, cells were incubated with 100 μL of fresh medium for 5 h. Cells were then added with 10 μL of alamar blue (YEASEN) per well. Cells were cultivated 24 h and the fluorescence of each well was measured with an excitation wavelength at 530 nm and an emission wavelength at 590 nm using BioTek plate reader (BioTek). The percentage of cell survival in the treated wells was calculated relative to the control-treated wells.

### Generation of rAAV in HEK-293T packaging cells

HEK-293T cells were seeded in the 75-cm^2^ flask (5×10^6^ cells/flask) and cultivated for 10‒16 h. Then the cells were cotransfected with 4 μg of plasmid pAAV-RC (Stratagene), 4 μg of plasmid pHelper (Stratagene), and 4 μg of one of following plasmids, including pAAV-HOsite-TMCP, pAAV-TsgRNA-dCas9-VP64, pAAV-HO, and pAAV-MCS (Stratagene), by using Lipofectamine 2000 (ThermoFisher Scientific) according to manufacturer’s instruction, respectively. The transfected cells were incubated in 37 °C and 5% CO_2_ for 72 h. The cells were scraped off and subjected three rounds of freeze–thaw–vortex cycles to release rAAV. The virus was purified with the PEG 8000 (Sigma) method and named as rAAV-HOsite-TMCP, rAAV-TsgRNA-dCas9-VP64, and rAAV-HO, rAAV-MCS, respectively.

### Measurement of virus titre

The titration of rAAV was analyzed by quantitative PCR (qPCR), and performed using the ABI StepOne Plus real-time PCR system (Applied Biosystems) with primers HO-TMCP q-F, HO-TMCP q-R, TsgRNA-dCas9 q-F, TsgRNA-dCas9 q-R, HO q-F and HO q-R. The PCR reaction mixture was performed in 20 μL of final volume using the 10 μL of SYBR Green Real-time PCR Master Mix (2×) (Roche) supplemented with 0.25 µM of primers, and 2 μL of twice serially diluted template DNA according to the manufacturer’s instructions. The primers used in qPCR were provided in Table S6. Each sample and blank control were run in at least three replicates of reactions in a 96-well optical plate. The PCR profile contained an initial denaturation step at 95 °C for 10 min followed by 45 cycles of 95 °C for 15 s and 60 °C for 1 min, followed by a melt curve stage of 95 °C for 15 s, 60 °C for 1 min and 95 °C for 15 s. Data analysis was performed using the Applied Biosystems StepOne software v2.3.

### Test of rAAVs with in vitro cultivated cells

The HepG2, Hepa1-6, MRC-5, and HL7702 cells were seeded into 24-well plates (0.5×10^5^ cells/well) and cultivated for 12‒20 h. Cells were then transfected with the viruses including rAAV-HOsite-TMCP, rAAV-TsgRNA-dCas9-VP64, rAAV-HO, and rAAV-MCS at the dose of 1×10^4^ vg per cell. The transfected cells were incubated at 37 °C and 5% CO_2_ for another 24 h. Cells were stained with Acridine orange. Cells were washed twice with PBS, then stained with 100 μg/mL acridine orange (Solarbio) for 10 min at room temperature. All cells were then observed and photographed under a fluorescence microscope (Olympus) at a constant magnification of 200×.

### Animal experiment

Four-week-old female nude mice purchased from the Model Animal Research Center of Nanjing (Nanjing, China) were used as the experimental animals. Each mouse weighed 18-22 g and was maintained under specific pathogen-free conditions. All animal experiments in this study followed the guidelines and ethics of the Animal Care and Use Committee of Southeast University (Nanjing, China).

In the first animal experiment, the nude mice were subcutaneously transplanted with 1×10^7^ of Hepa1-6 cells mixed with 1×10^9^ vg of rAAV-MCS or rAAV-TSD (an equivalent mixture of three viruses including rAAV-HO, rAAV-TsgRNA-dCas9-VP64, and rAAV-HOsite-TMCP), respectively. After two weeks, all mice were sacrificed and photographed. The tumor size was measured.

In the second animal experiment, the nude mice subcutaneously injected with 1×10^7^ Hepa1-6 cells to produce the tumor-burdened mice. After one week, the tumor mice were intravenously injected with 1×10^9^ vg of purified rAAV-TSD and rAAV-MCS, respectively. After one week, all mice were sacrificed and photographed. The tumor size was measured.

### Detection of virus presence in tissues with qPCR

The heart, liver, spleen, lung, kidney and tumor tissues were collected from mice in the second animal experiment. The gDNA was extracted from above tissues using TIANamp Genomic DNA Kit (TIANGEN) according to the manufacturer’s instructions. The AAV DNA abundance were analyzed by quantitative PCR (qPCR) with primers MCS-qF and MCS-qR using the ABI StepOne Plus real-time PCR system (Applied Biosystems). The real-time PCR reaction mixture was performed in a total volume of 20 µL, containing 10 µL of SYBR Green Real-time PCR Master Mix (2×) (Roche), 0.25 µM of primers, and 10 ng of gDNA. Real-time PCR profile contained an initial denaturation step at 95 °C for 10 min, followed by 45 cycles of 95 °C for 15 s and 60 °C for 1 min, then followed by a melt curve stage of 95 °C for 15 s, 60 °C for 1 min and 95 °C for 15 s. All real-time PCR detections and blank control were performed in at least three technical replicates in a 96-well optical plate. Melting curve analysis revealed a single PCR product. The threshold value Ct for each individual PCR product was calculated by the Applied Biosystems StepOne software v2.3, and Ct values were normalized by subtracting the Ct values obtained for GAPDH (GAPDH-qF1 and GAPDH-qR1). The primers used in real-time PCR were provided in Table S6. The resulting ΔCt values were then used to calculate relative abundance of viral DNA as relative quantity (RQ) according to the equation: RQ= 2^−ΔΔCt^.

### Detection of effector gene expression in tissues with qPCR

The heart, liver, spleen, lung, kidney and tumor tissues were randomly collected from mice in the second animal experiment. Total RNA was extracted from above tissues with Trizol^®^ reagent (Invitrogen), then 500 ng of total RNA was reversely transcribed into cDNA using PrimeScript^TM^ RT Master Mix (TaKaRa) according to the manufacturer’s instructions in a total volume of 10 µL. The Cas9 mRNA expression levels were analyzed by quantitative PCR (qPCR) with primers Cas9-qF and Cas9-qR using the ABI StepOne Plus real-time PCR system (Applied Biosystems). The real-time PCR reaction mixture was performed in a total volume of 20 µL, containing 10 µL of 2× SYBR Green Real-time PCR Master Mix (Roche), 0.25 µM of primers, and 2 µL cDNA. Real-time PCR profile contained an initial denaturation step at 95 °C for 10 min, followed by 45 cycles of 95 °C for 15 s and 60 °C for 1 min, then followed by a melt curve stage of 95 °C for 15 s, 60 °C for 1 min and 95 °C for 15 s. All real-time PCR detections and blank control were performed in at least three technical replicates in a 96-well optical plate. Melting curve analysis revealed a single PCR product. The threshold value Ct for each individual PCR product was calculated by the Applied Biosystems StepOne software v2.3, and Ct values were normalized by subtracting the Ct values obtained for *GAPDH* (GAPDH-qF2 and GAPDH-qR2). The primers used in real-time PCR were provided in Table S6. The resulting ΔCt values were then used to calculate relative changes of Cas9 mRNA expression as relative quantity (RQ) according to the equation: RQ= 2^−ΔΔCt^.

### Test the Tage system with TALE as transcription factor

293T cells were transfected with various combinations of vectors including sTMEP, TALE, and C1-EGFP. After transfection, cells were cultivated for 24 h. Cells were then observed and photographed with a fluorescence microscope (Olympus) at a constant magnification of 200×. The fluorescence intensity of cells were quantified with a flow cytometry (Calibur, BD, USA) and the mean fluorescent intensity (MFI) was calculated by BD software of flow cytometry.

## Statistical analyses

Data were expressed as means values±standard deviation (SD) at least three independent biological or experimental replicated, and analyzed by a two-tailed unpaired Student’s *t*-test. Differences at *p < 0.05* were considered statistical significance.

---

**Fig.S1.**
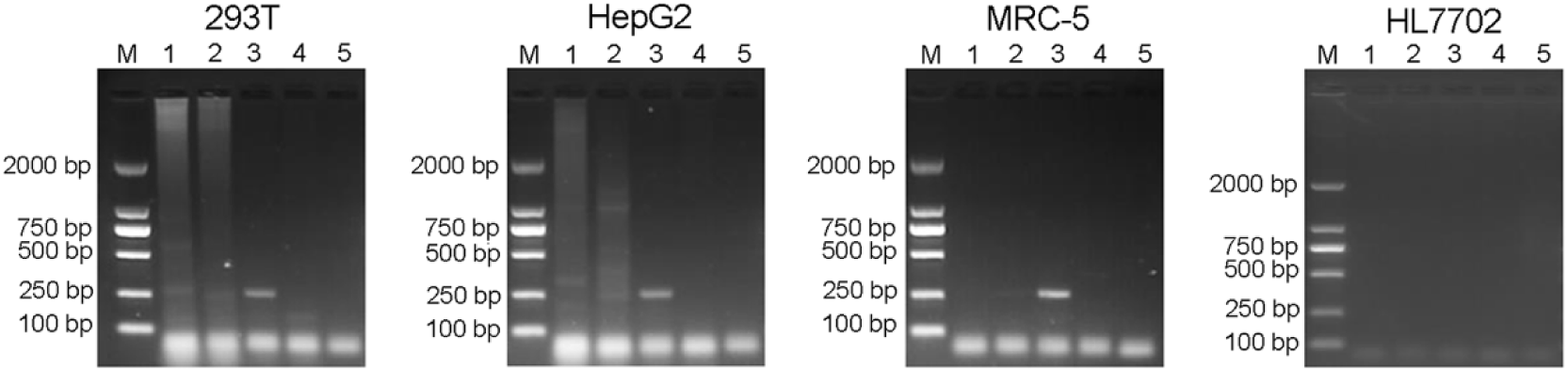
Evaluation the specificity of Tage system in different cell lines. PCR am plification assay of telomerase elongation products of 293T, HepG2, MEC-5 and HL7702 cell lines. M, DL2000 DNA marker; Lane 1, cells co-transfected with sTMEP & TsgRNA & dCas9-VP64; Lane 2, cells transfected with sTMEP; Lane 3, cells transfected with bTMEP; Lane 4, cells transfected with lipofectin; Lane 5, PCR negative control.

**Fig.S2.**
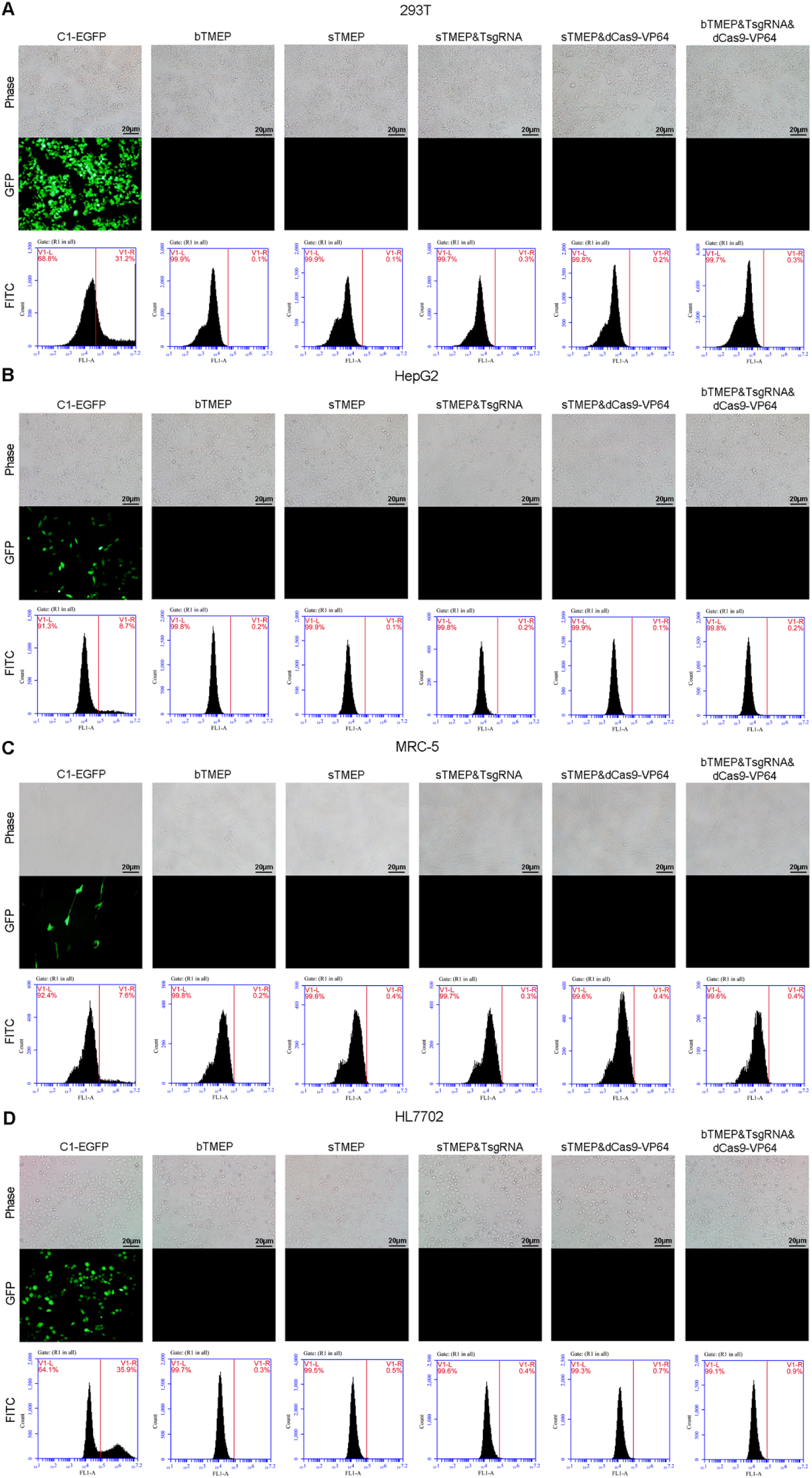
Evaluation the specificity of Tage system in different cell lines. (A) The fluorescence microscope pictures and representative flow cytometry analysis of Tage system specificity of control groups in 293T cell lines. (B) The fluorescence microscope pictures and representative flow cytometry analysis of Tage system specificity of control groups in HepG2 cell lines. (C) The fluorescence microscope pictures and representative flow cytometry analysis of Tage system specificity of control groups in MRC-5 cell lines. (D) The fluorescence microscope pictures and representative flow cytometry analysis of Tage system specificity of control groups in HL7702 cell lines.

**Fig.S3.**
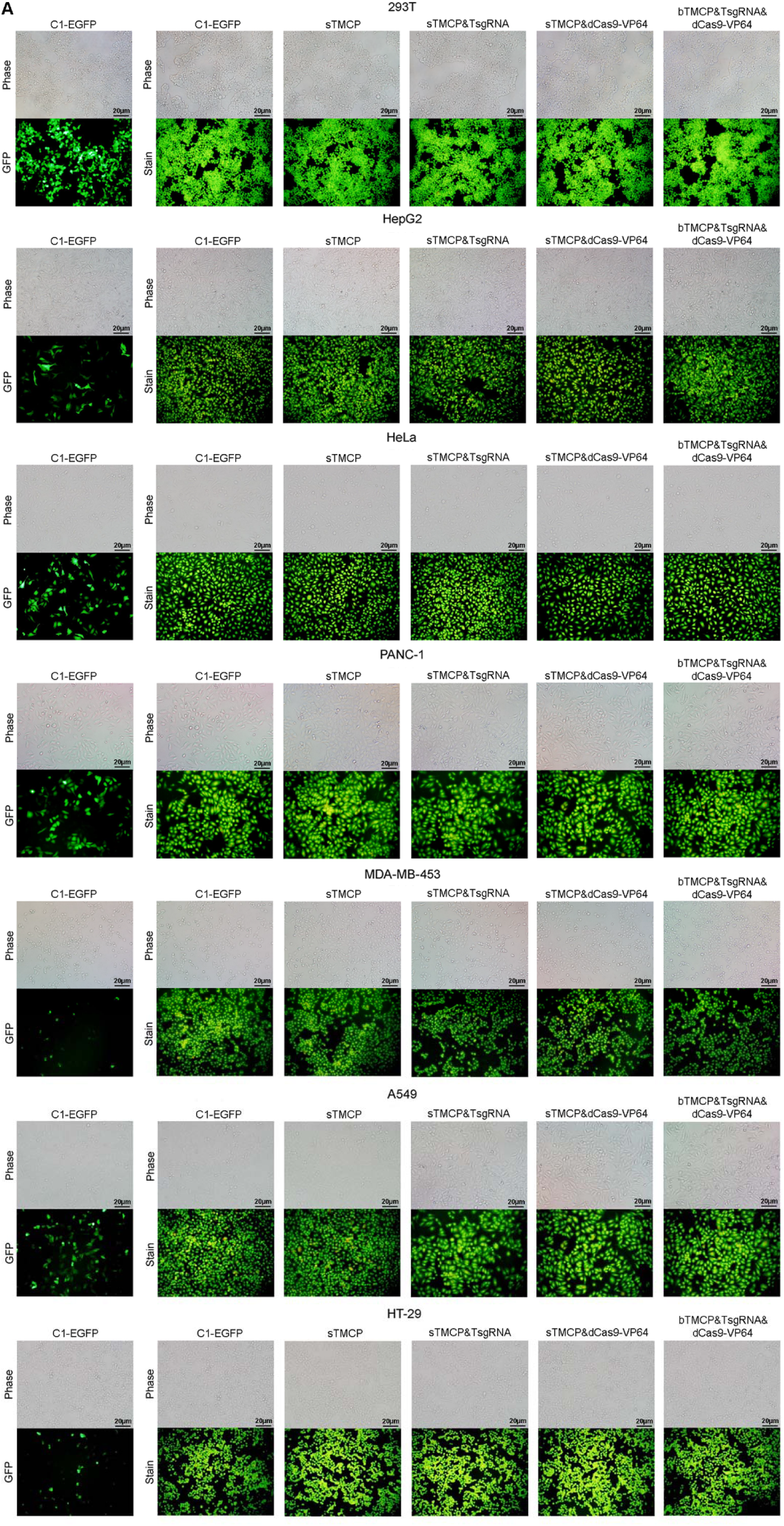
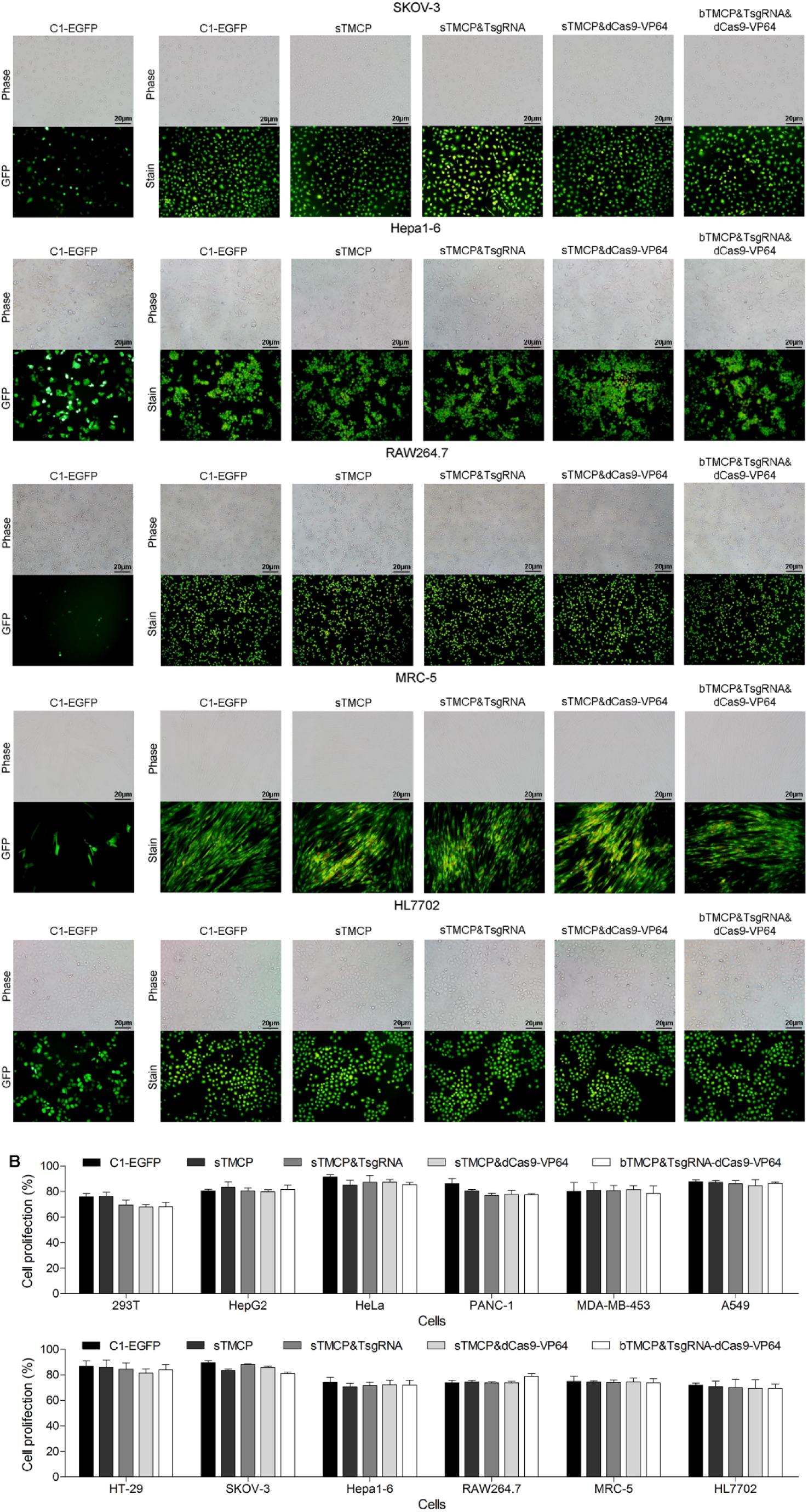
Kill tumor cells with the Tage system. (A) The microscope pictures and representative acridine orange staining pictures of control groups of Tage system to kill tumor cells in 293T, HepG2, HeLa, PANC-1, MDA-MB-453, A549, HT-29, SKOV-3, Hepa1-6, RAW264.7, MRC-5 and HL7702 cell lines. (B) The alamar blue analysis control groups of Tage system to kill tumor cells in 293T, HepG2, HeLa, PANC-1, MDA-MB-453, A549, HT-29, SKOV-3, Hepa1-6, RAW264.7, MRC-5 and HL7702 cell lines.

**Fig.S4.**
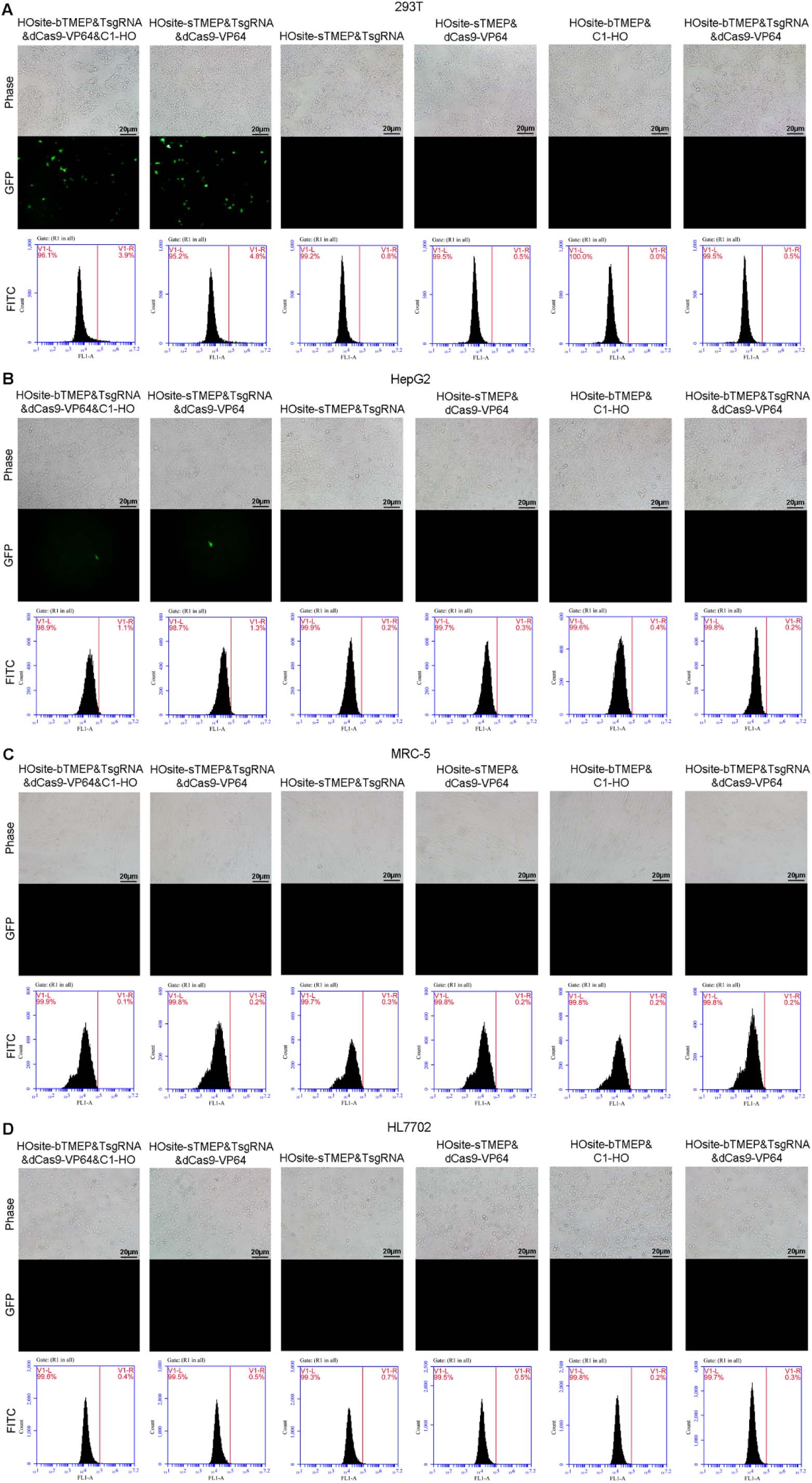
Evaluation of elongation of 4 bases 3ʹ single-stranded stick-ended fragments in different cell lines. (A) The fluorescence microscope pictures of 293T cell lines. (B) The fluorescence microscope pictures of HepG2 cell lines. (C) The fluorescence microscope pictures of MRC-5 cell lines. (D) The fluorescence microscope pictures of HL7702 cell lines.

**Fig.S5.**
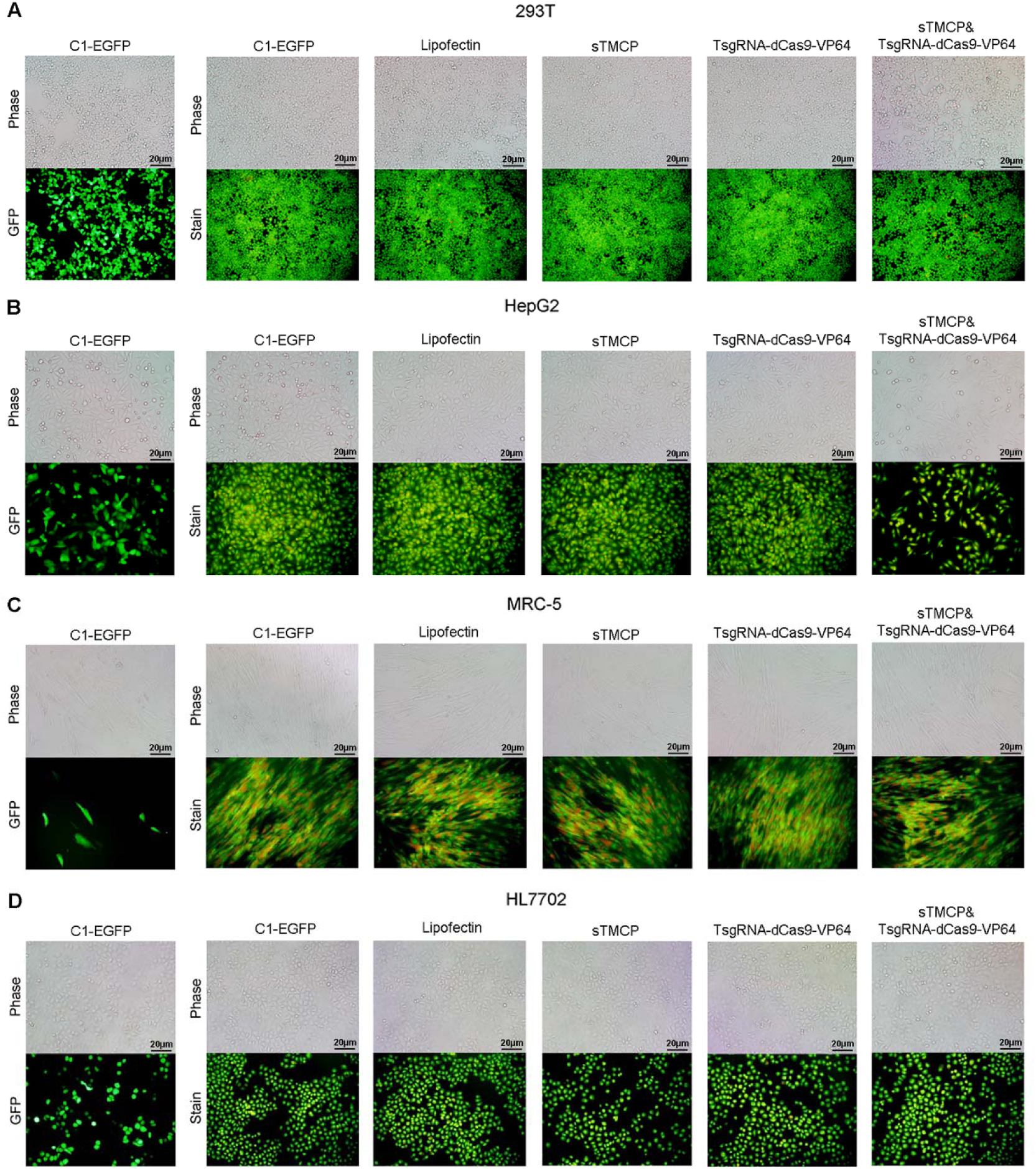
Application of TsgRNA fused with dCas9-VP64 of Tage system to kill tumor cells. (A) The microscope pictures and representative acridine orange staining pictures of 293T cell lines. (B) The microscope pictures and representative acridine orange staining pictures of HepG2 cell lines. (C) The microscope pictures and representative acridine orange staining pictures of MRC-5 cell lines. (D) The microscope pictures and representative acridine orange staining pictures of HL7702 cell lines.

**Fig.S6.**
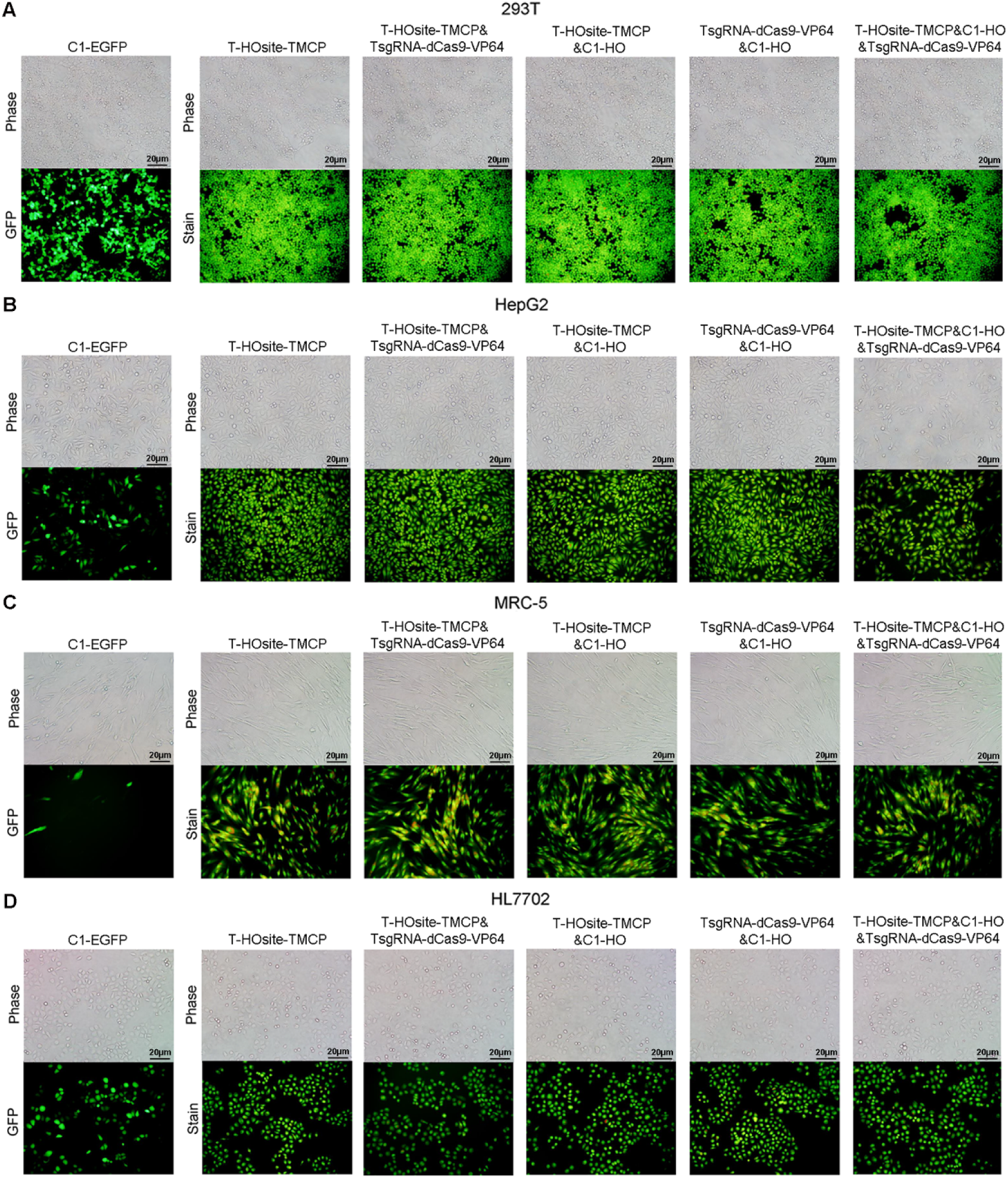
Combined TsgRNA-dCas9-VP64 with HO enzymes of Tage system to kill tumor cells. (A) The microscope pictures and representative acridine orange staining pictures of 293T cell lines. (B) The microscope pictures and representative acridine orange staining pictures of HepG2 cell lines. (C) The microscope pictures and representative acridine orange staining pictures of MRC-5 cell lines. (D) The microscope pictures and representative acridine orange staining pictures of HL7702 cell lines.

**Fig.S7.**
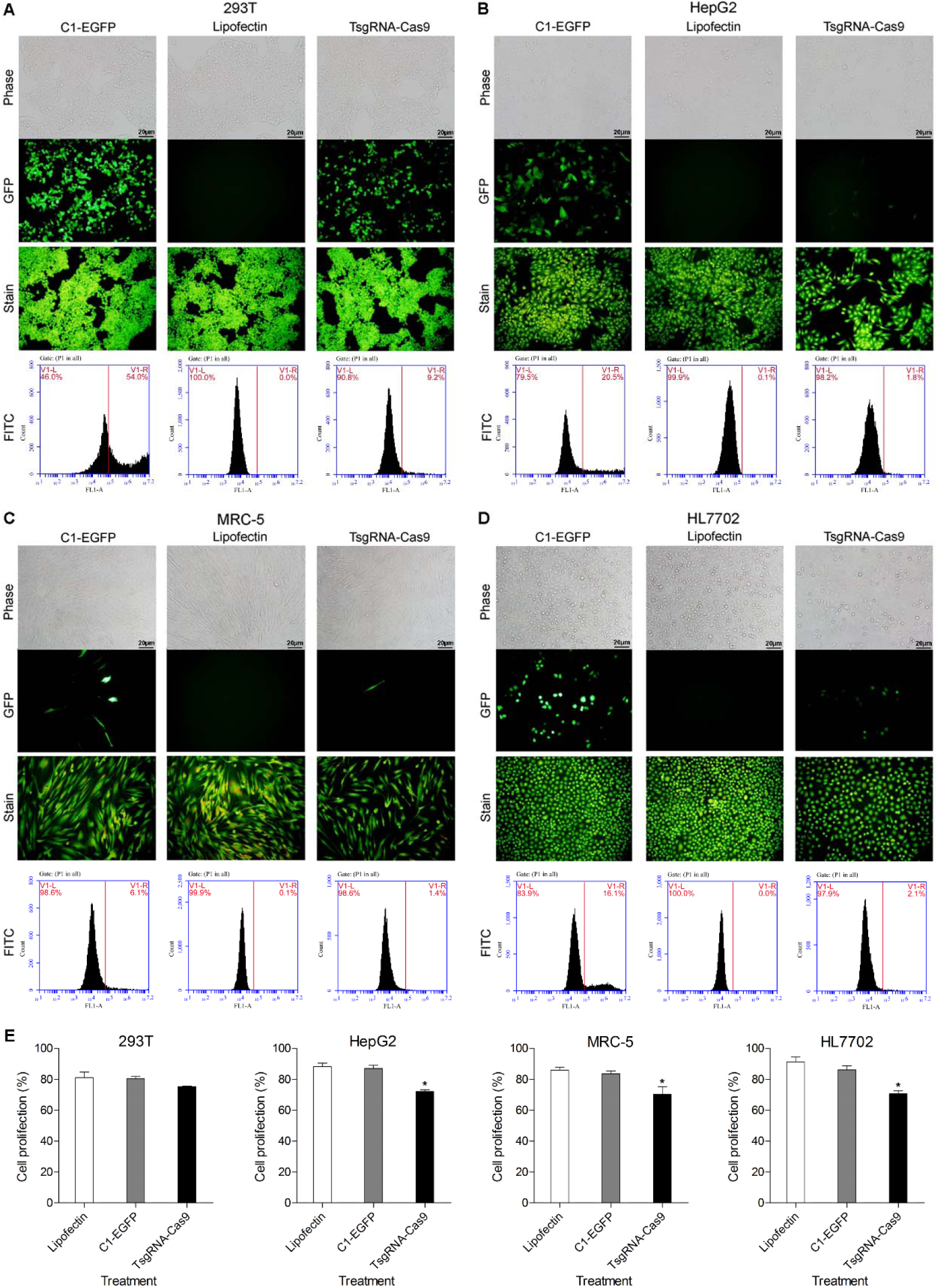
Directly cleavage of telomere DNA with TsgRNA-Cas9 vector in different cell lines. (A) The fluorescence microscope pictures, acridine orange staining pictures and representative flow cytometry analysis of 293T cell lines. (B) The fluorescence microscope pictures, acridine orange staining pictures and representative flow cytometry analysis of HepG2 cell lines. (C) The fluorescence microscope pictures, acridine orange staining pictures and representative flow cytometry analysis of MRC-5 cell lines. (D) The fluorescence microscope pictures, acridine orange staining pictures and representative flow cytometry analysis of HL7702 cell lines. (E) The alamar blue analysis of 293T, HepG2, MRC-5 and HL7702 cell lines.

**Fig.S8.**
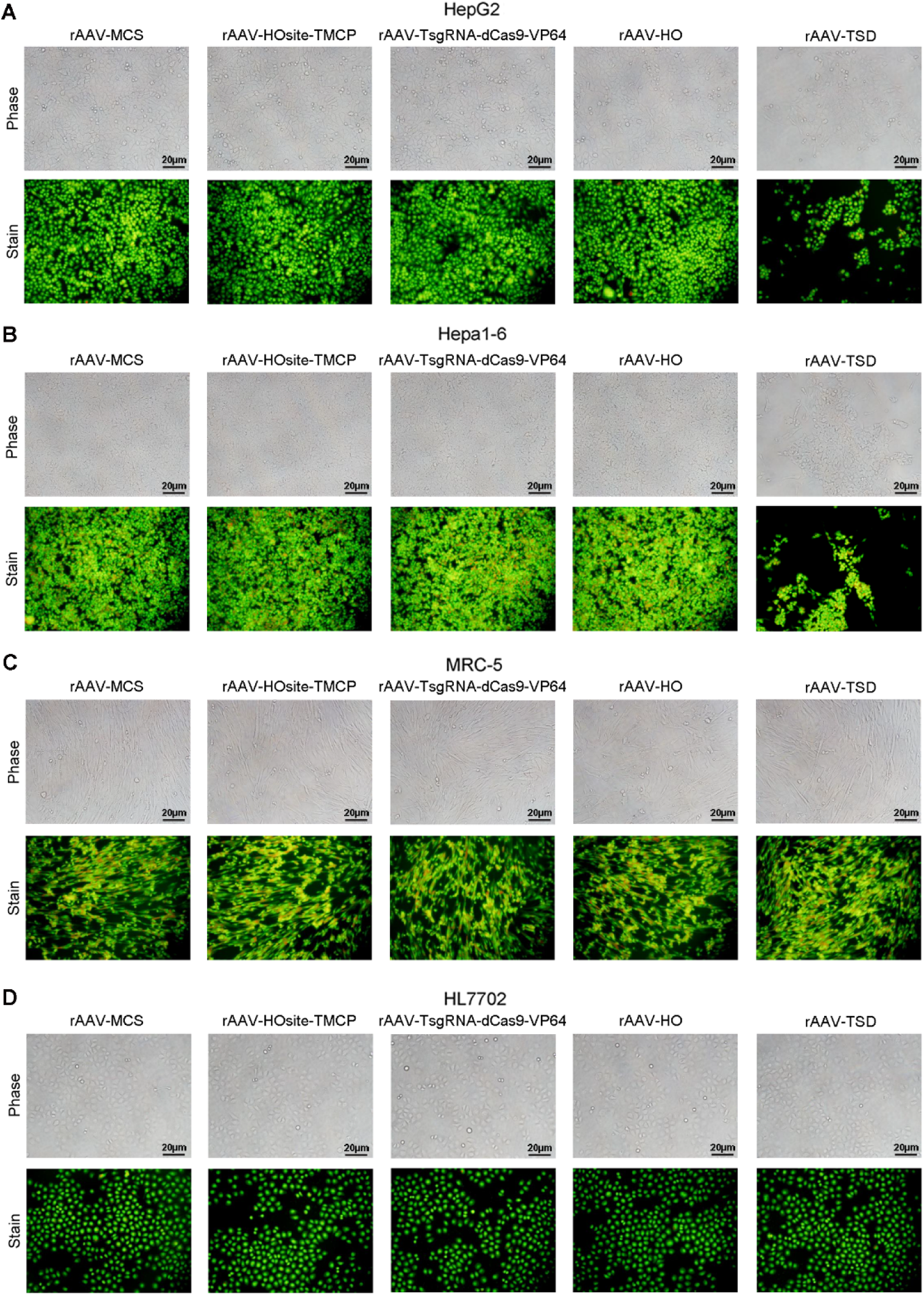
Evaluation of rAAV to kill tumor cells. (A) The microscope pictures and representative acridine orange staining pictures of HepG2 cell lines. (B) The microscope pictures and representative acridine orange staining pictures of Hepa1-6 cell lines. (C) The microscope pictures and representative acridine orange staining pictures of MRC-5 cell lines. (D) The microscope pictures and representative acridine orange staining pictures of HL7702 cell lines. rAAV-TSD refers to the equivalent mixture of three viruses including rAAV-HOsite-TMCP & rAAV-TsgRNA-dCas9-VP64 & rAAV-HO.

**Fig.S9.**
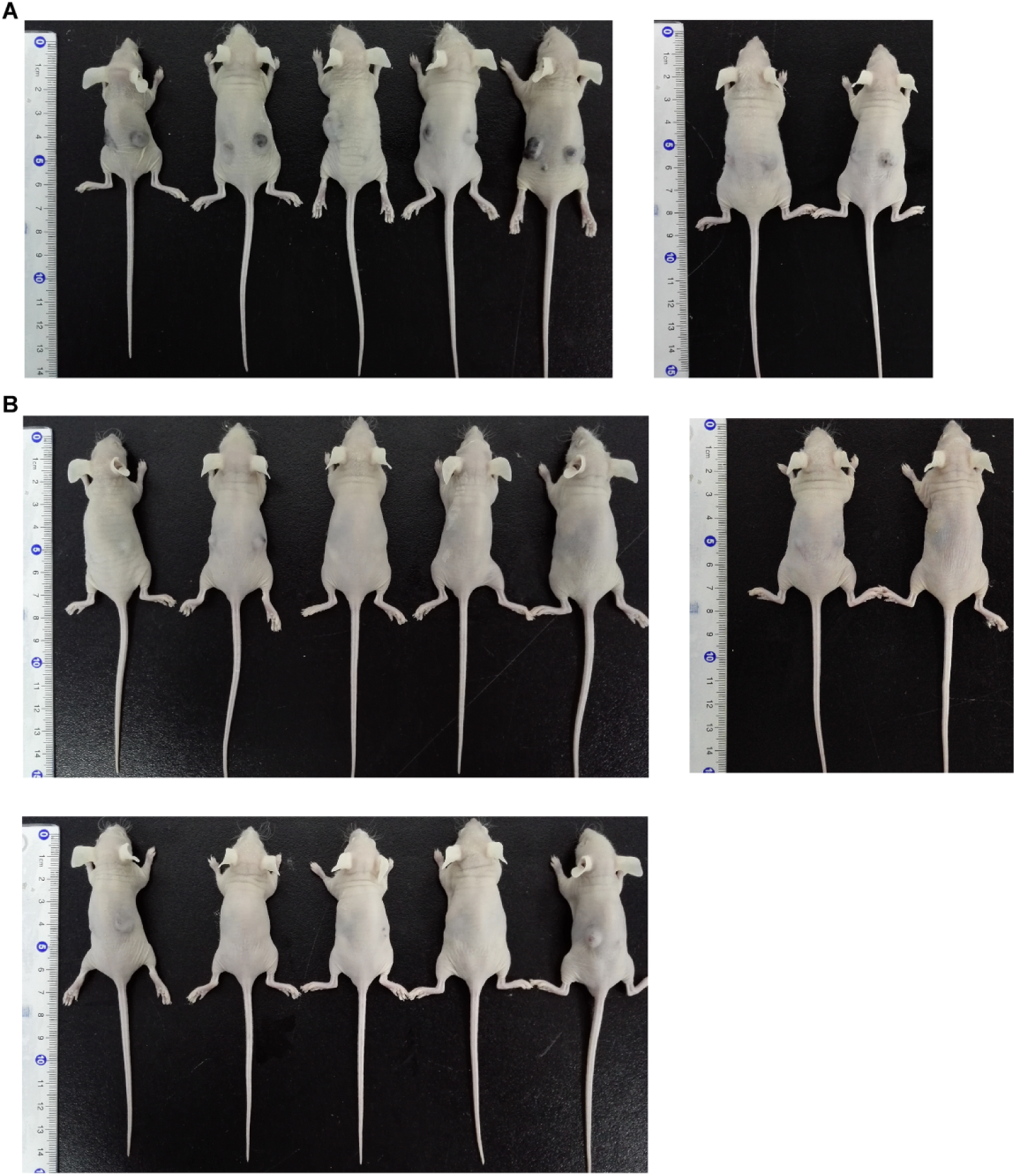
Application Tage system for cancer therapy *in vivo* via subcutaneously injected with rAAV. (A) Mice subcutaneously transplanted with the Hepa1-6 cells mixed with rAAV-MCS. (B) Mice subcutaneously transplanted with the Hepa1-6 cells mixed with rAAV-TSD. rAAV-TSD refers to the equivalent mixture of three rAVVs, including rAAV-HOsite-TMCP, rAAV-TsgRNA-dCas9-VP64, and rAAV-HO.

**Fig.S10.**
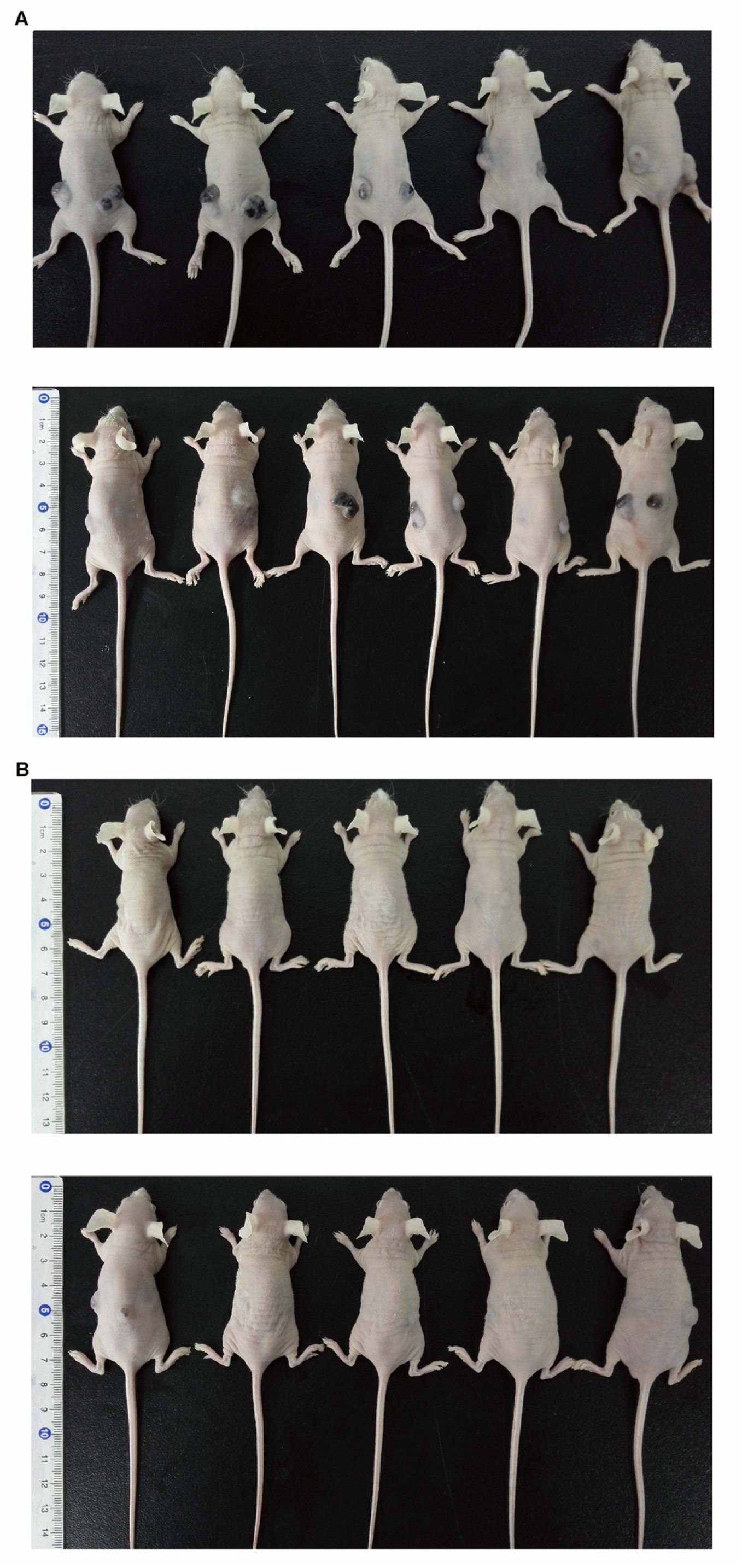
Application of Tage system to cancer therapy *in vivo* by intravenously injecting rAAV into the mice with tumors. (A) Mice intravenously injected with rAAV-MCS. (B) Mice intravenously injected with rAAV-TSD. rAAV-TSD refers to the equivalent mixture of three rAVVs, including rAAV-HOsite-TMCP, rAAV-TsgRNA-dCas9-VP64, and rAAV-HO.

**Fig.S11.**
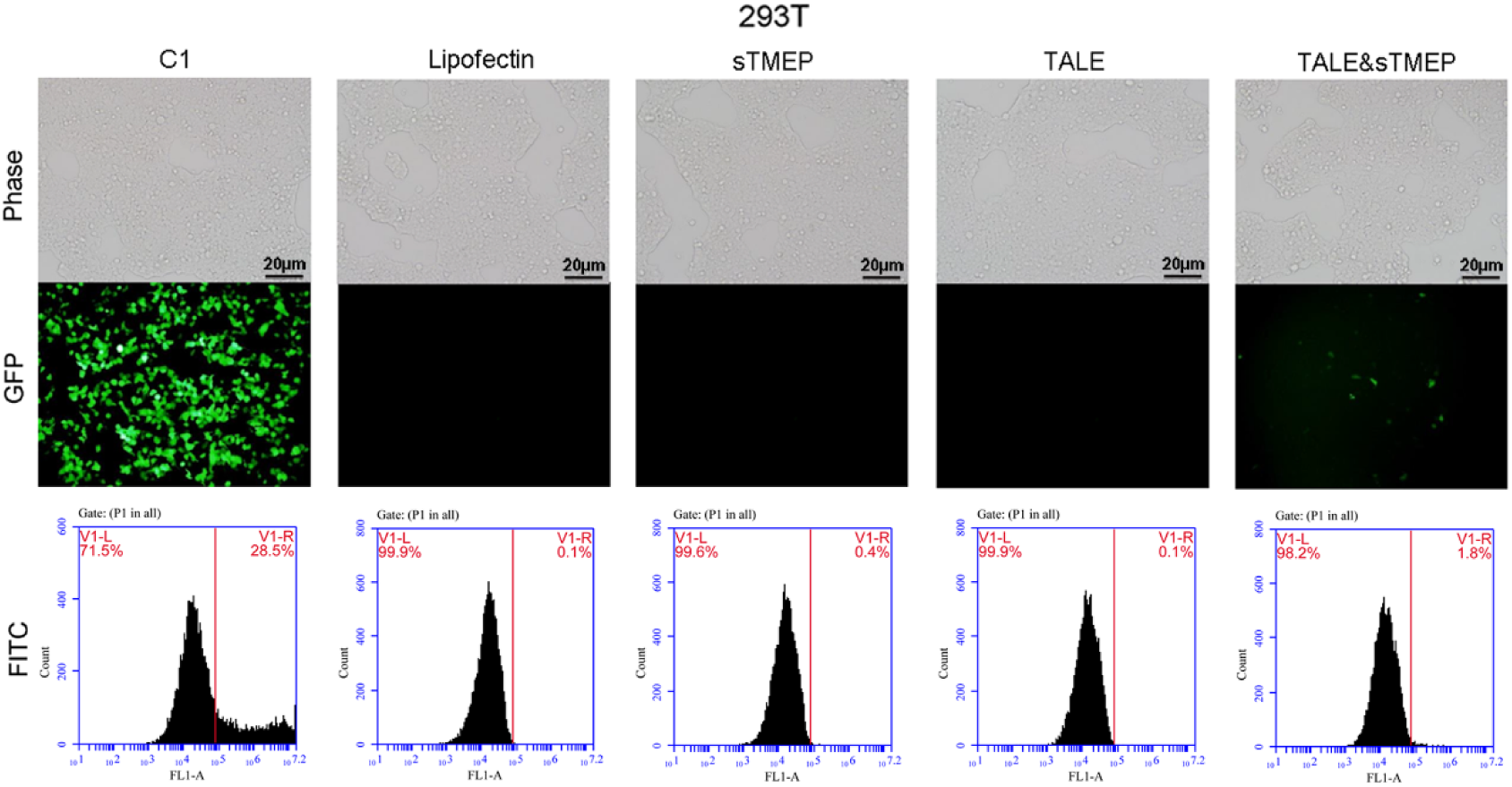
Activation of ZsGreen expression by the Tage system using TALE as artificial transcription factor. 293T cell was co-transfected with sTMEP & TALE. Cell was photographed with microscope and the fluorescence was quantified with flow cytometry. Cell was also transfected by the sole sTMEP and TALE as negative controls. Cell transfected with C1-EGFP was used as a control for monitoring transfection.

**Table S1.**
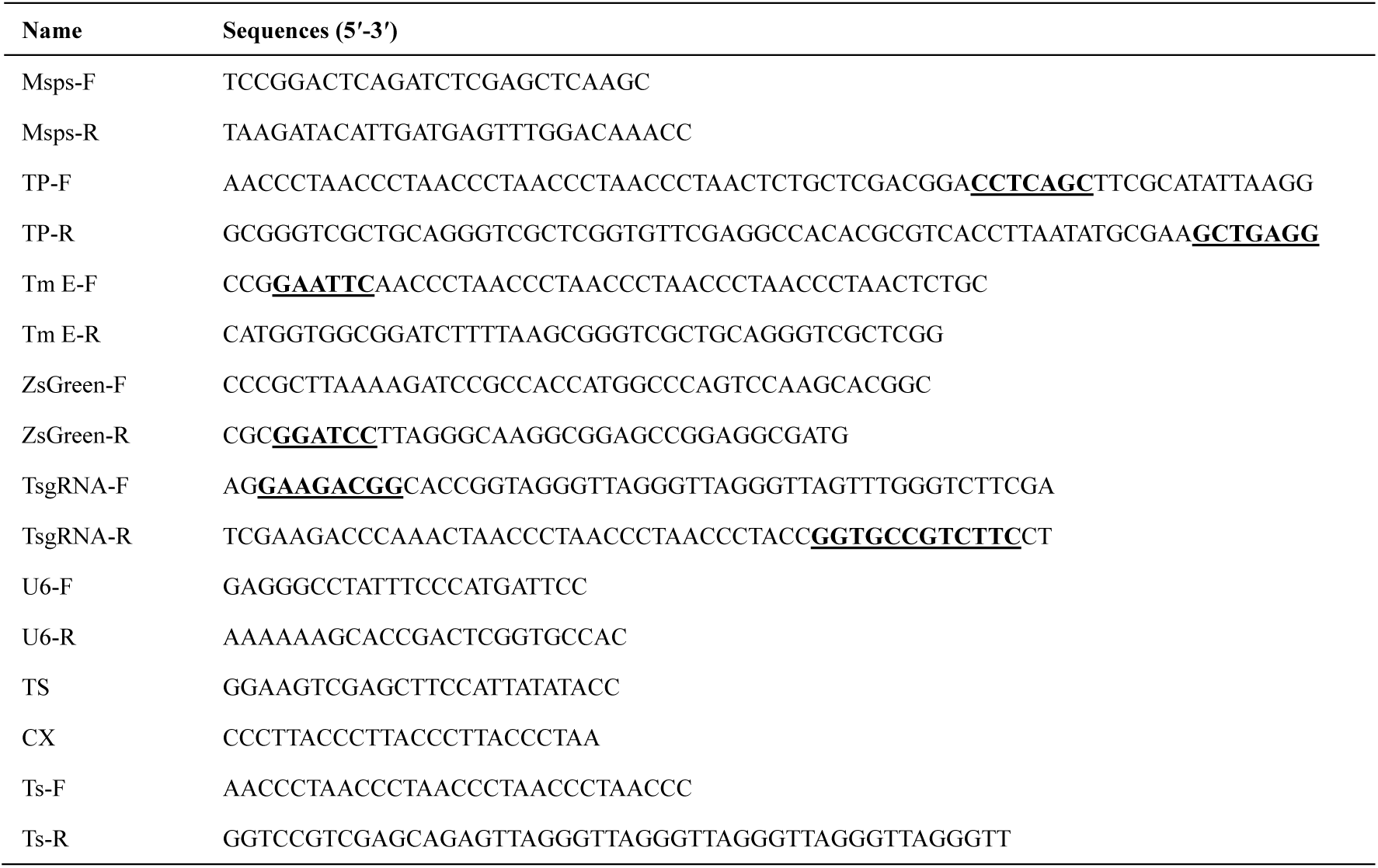
Oligonucleotides used in TMEP experiment. The bold and underline indicated the digest sites.

**Table S2.**
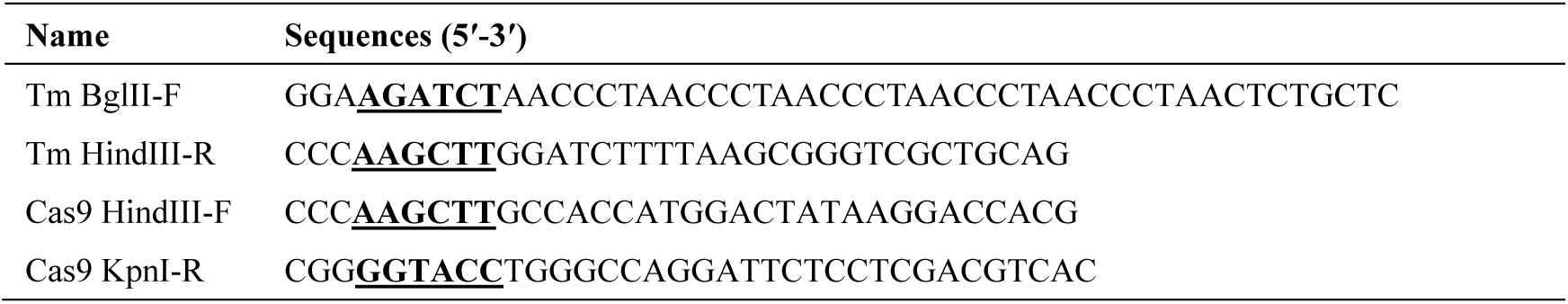
Oligonucleotides used in TMCP experiment. The bold and underline indicated the digest sites.

**Table S3.**
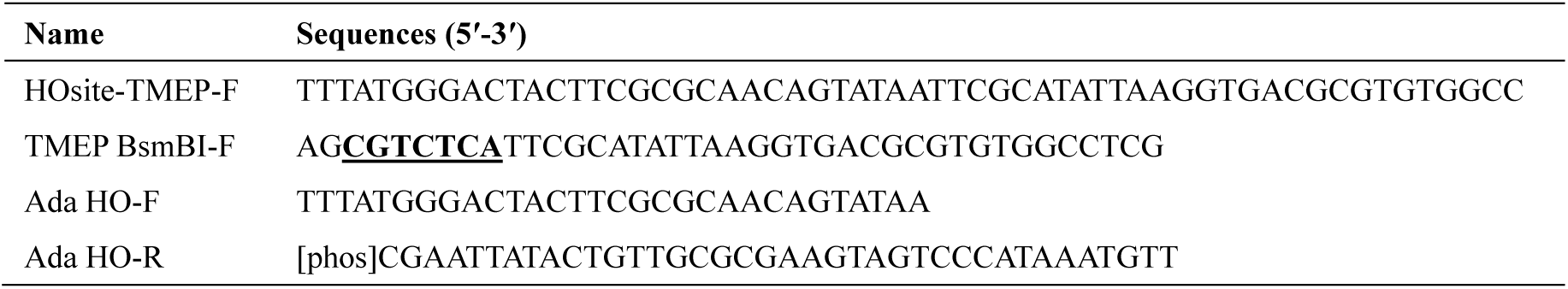
Oligonucleotides used in HOsite-TMEP experiment. The bold and underline indicate the digestion sites.

**Table S4.**
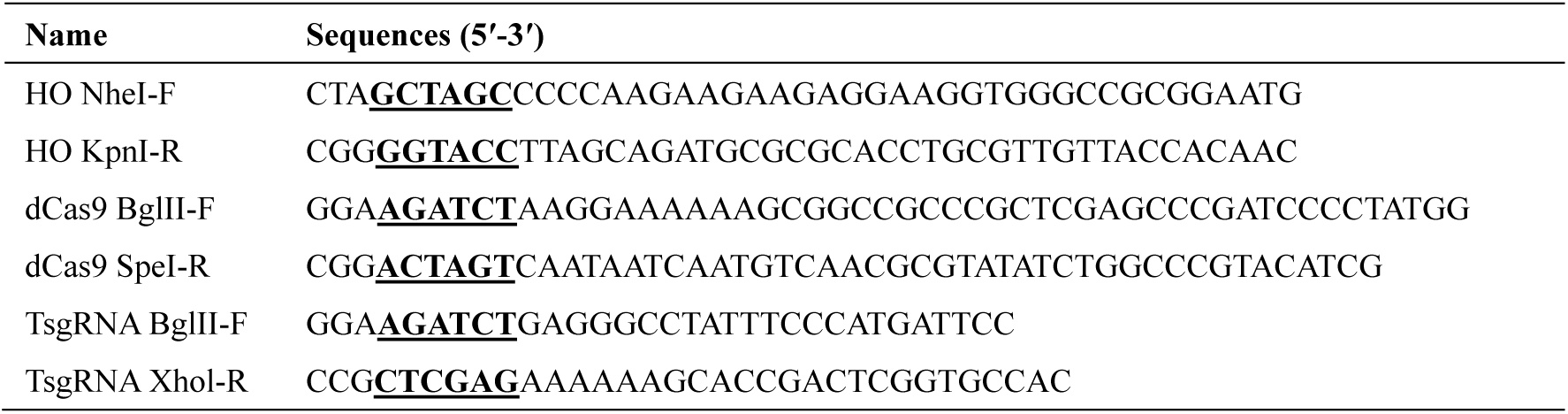
Oligonucleotides used in HOsite-TMCP experiment. The bold and underline indicated the digest sites.

**Table S5.**
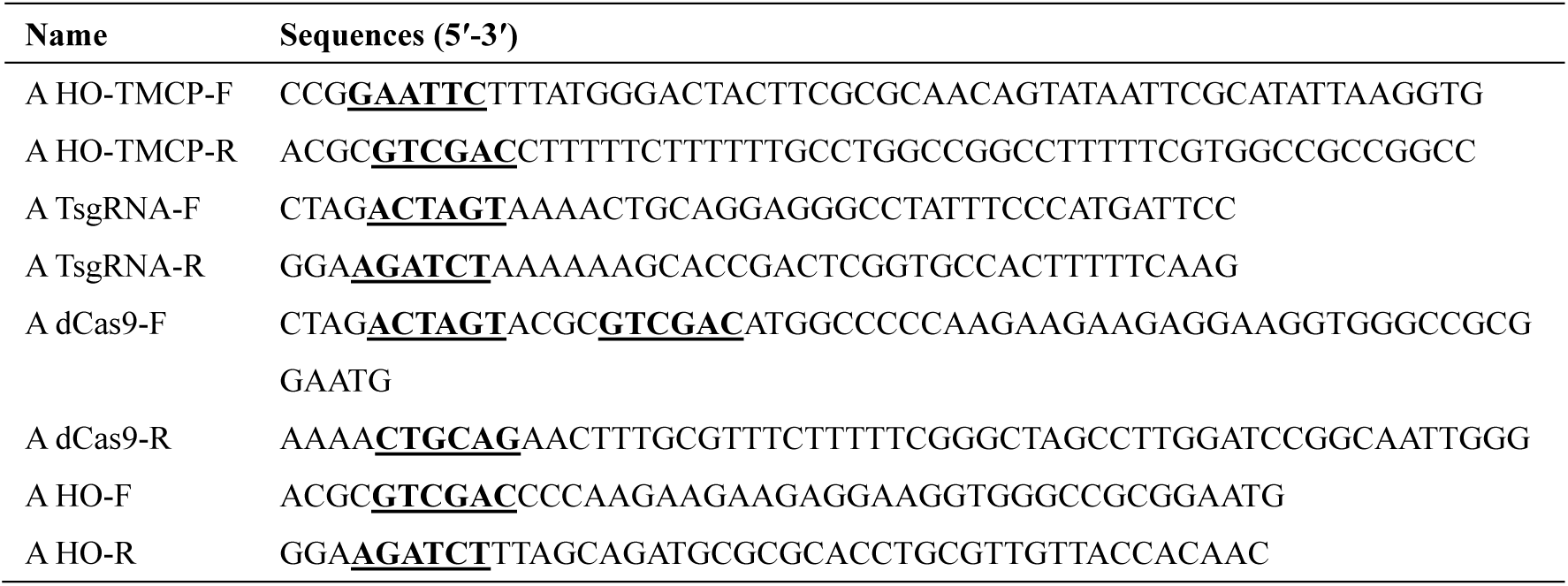
Oligonucleotides used in rAAV experiment. The bold and underline indicate the digestion sites.

**Table S6.**
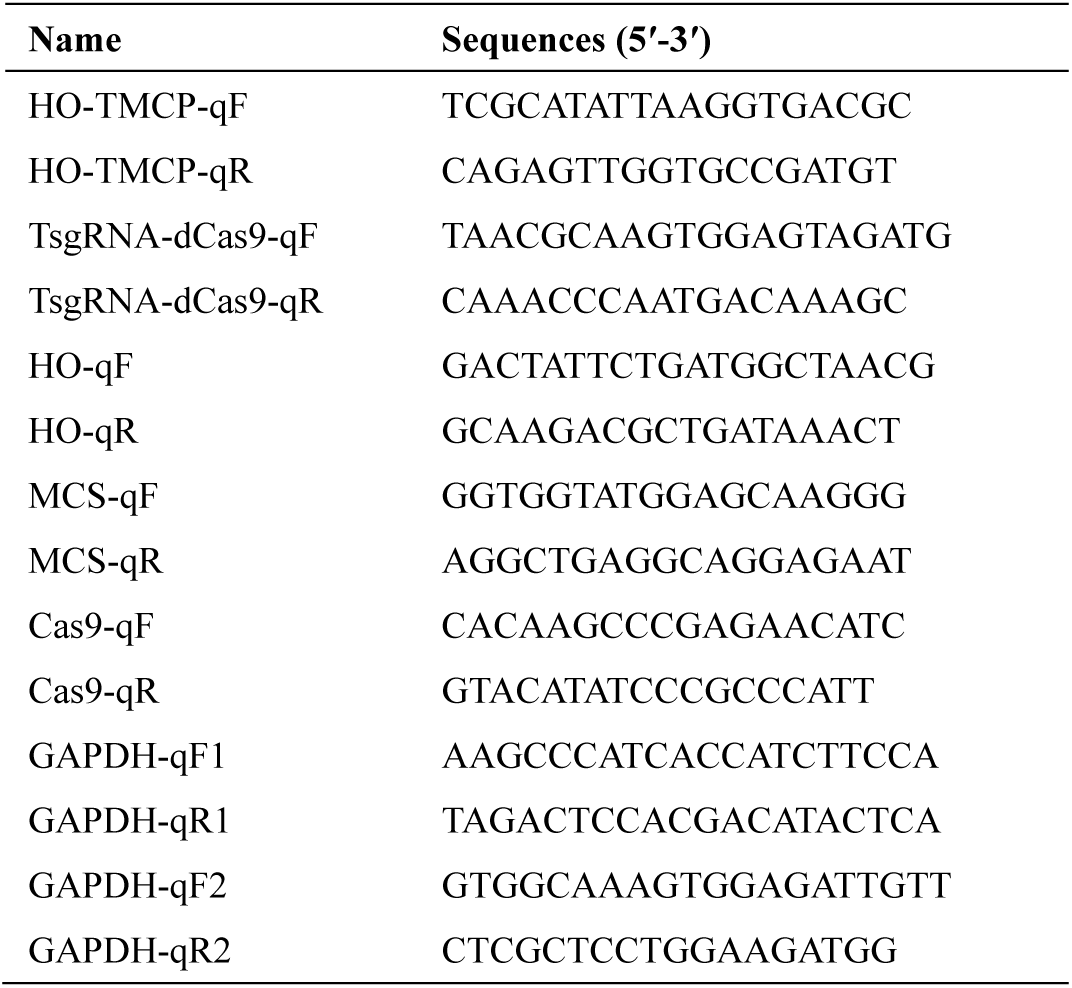
Oligonucleotides used in qPCR.

## Supplementary sequences

### >TMEP: Telomere repeat sequences + Mini-TK promoter + ZsGreen + SV40 poly(A) signal

AACCCTAACCCTAACCCTAACCCTAACCCTAACTCTGCTCGACGGACC^^^TCAGCTTC GCATATTAAGGTGACGCGTGTGGCCTCGAACACCGAGCGACCCTGCAGCGACCCGCTTAAAAGATCCGCCACCATGGCCCAGTCCAAGCACGGCCTGACCAAGGAGATGACCATGAAGTACCGCATGGAGGGCTGCGTGGACGGCCACAAGTTCGTGATCACCGGCGAGGGCATCGGCTACCCCTTCAAGGGCAAGCAGGCCATCAACCTGTGCGTGGTGGAGGGCGGCCCCTTGCCCTTCGCCGAGGACATCTTGTCCGCCGCCTTCATGTACGGCAACCGCGTGTTCACCGAGTACCCCCAGGACATCGTCGACTACTTCAAGAACTCCTGCCCCGCCGGCTACACCTGGGACCGCTCCTTCCTGTTCGAGGACGGCGCCGTGTGCATCTGCAACGCCGACATCACCGTGAGCGTGGAGGAGAACTGCATGTACCACGAGTCCAAGTTCTACGGCGTGAACTTCCCCGCCGACGGCCCCGTGATGAAGAAGATGACCGACAACTGGGAGCCCTCCTGCGAGAAGATCATCCCCGTGCCCAAGCAGGGCATCTTGAAGGGCGACGTGAGCATGTACCTGCTGCTGAAGGACGGTGGCCGCTTGCGCTGCCAGTTCGACACCGTGTACAAGGCCAAGTCCGTGCCCCGCAAGATGCCCGACTGGCACTTCATCCAGCACAAGCTGACCCGCGAGGACCGCAGCGACGCCAAGAACCAGAAGTGGCACCTGACCGAGCACGCCATCGCCTCCGGCTCCGCCTTGCCCTAAGGATCCACCGGATCTAGATAACTGATCATAATCAGCCATACCACATTTGTAGAGGTTTTACTTGCTTTAAAAAACCTCCCACACCTCCCCCTGAACCTGAAACATAAAATGAATGCAATTGTTGTTGTTAACTTGTTTATTGCAGCTTATAATGGTTACAAATAAAGCAATAGCATCACAAATTTCACAAATAAAGCATTTTTTTCACTGCATTCTAGTTGTGGTTTGTCCAAACTCATCAATGTATCTTA

### >TsgRNA: U6 promoter + telomere target sequence + gRNA scaffold

GAGGGCCTATTTCCCATGATTCCTTCATATTTGCATATACGATACAAGGCTGTTAGAGAGATAATTGGAATTAATTTGACTGTAAACACAAAGATATTAGTACAAAATACGTGACGTAGAAAGTAATAATTTCTTGGGTAGTTTGCAGTTTTAAAATTATGTTTTAAAATGGACTATCATATGCTTACCGTAACTTGAAAGTATTTCGATTTCTTGGCTTTATATATCTTGTGGAAAGGACGAAACACCGTAGGGTTAGGGTTAGGGTTAGTTTTAGAGCTAGAAATAGCAAGTTAAAATAAGGCTAGTCCGTTATCAACTTGAAAAAGTGGCACCGAGTCGGTGCTTTTTT

### >TMCP: Telomere repeat sequences + Mini-TK promoter + SV40 NLS + Cas9 + SV40 NLS + SV40 poly(A) signal

AACCCTAACCCTAACCCTAACCCTAACCCTAACTCTGCTCGACGGACC^TCAGCTT CGCATATTAAGGTGACGCGTGTGGCCTCGAACACCGAGCGACCCTGCAGCGACCC GCTTAAAAGATCCAAGCTTGCCACCATGGACTATAAGGACCACGACGGAGACTAC AAGGATCATGATATTGATTACAAAGACGATGACGATAAGATGGCCCCAAAGAAGAA GCGGAAGGTCGGTATCCACGGAGTCCCAGCAGCCGACAAGAAGTACAGCATCGG CCTGGACATCGGCACCAACTCTGTGGGCTGGGCCGTGATCACCGACGAGTACAAG GTGCCCAGCAAGAAATTCAAGGTGCTGGGCAACACCGACCGGCACAGCATCAAG AAGAACCTGATCGGAGCCCTGCTGTTCGACAGCGGCGAAACAGCCGAGGCCACC CGGCTGAAGAGAACCGCCAGAAGAAGATACACCAGACGGAAGAACCGGATCTGC TATCTGCAAGAGATCTTCAGCAACGAGATGGCCAAGGTGGACGACAGCTTCTTCC ACAGACTGGAAGAGTCCTTCCTGGTGGAAGAGGATAAGAAGCACGAGCGGCACC CCATCTTCGGCAACATCGTGGACGAGGTGGCCTACCACGAGAAGTACCCCACCAT CTACCACCTGAGAAAGAAACTGGTGGACAGCACCGACAAGGCCGACCTGCGGCT GATCTATCTGGCCCTGGCCCACATGATCAAGTTCCGGGGCCACTTCCTGATCGAGG GCGACCTGAACCCCGACAACAGCGACGTGGACAAGCTGTTCATCCAGCTGGTGC AGACCTACAACCAGCTGTTCGAGGAAAACCCCATCAACGCCAGCGGCGTGGACG CCAAGGCCATCCTGTCTGCCAGACTGAGCAAGAGCAGACGGCTGGAAAATCTGAT CGCCCAGCTGCCCGGCGAGAAGAAGAATGGCCTGTTCGGAAACCTGATTGCCCTG AGCCTGGGCCTGACCCCCAACTTCAAGAGCAACTTCGACCTGGCCGAGGATGCCA AACTGCAGCTGAGCAAGGACACCTACGACGACGACCTGGACAACCTGCTGGCCC AGATCGGCGACCAGTACGCCGACCTGTTTCTGGCCGCCAAGAACCTGTCCGACGC CATCCTGCTGAGCGACATCCTGAGAGTGAACACCGAGATCACCAAGGCCCCCCTG AGCGCCTCTATGATCAAGAGATACGACGAGCACCACCAGGACCTGACCCTGCTGA AAGCTCTCGTGCGGCAGCAGCTGCCTGAGAAGTACAAAGAGATTTTCTTCGACCA GAGCAAGAACGGCTACGCCGGCTACATTGACGGCGGAGCCAGCCAGGAAGAGTT CTACAAGTTCATCAAGCCCATCCTGGAAAAGATGGACGGCACCGAGGAACTGCTC GTGAAGCTGAACAGAGAGGACCTGCTGCGGAAGCAGCGGACCTTCGACAACGGC AGCATCCCCCACCAGATCCACCTGGGAGAGCTGCACGCCATTCTGCGGCGGCAG GAAGATTTTTACCCATTCCTGAAGGACAACCGGGAAAAGATCGAGAAGATCCTGA CCTTCCGCATCCCCTACTACGTGGGCCCTCTGGCCAGGGGAAACAGCAGATTCGC CTGGATGACCAGAAAGAGCGAGGAAACCATCACCCCCTGGAACTTCGAGGAAGT GGTGGACAAGGGCGCTTCCGCCCAGAGCTTCATCGAGCGGATGACCAACTTCGAT AAGAACCTGCCCAACGAGAAGGTGCTGCCCAAGCACAGCCTGCTGTACGAGTACT TCACCGTGTATAACGAGCTGACCAAAGTGAAATACGTGACCGAGGGAATGAGAAA GCCCGCCTTCCTGAGCGGCGAGCAGAAAAAGGCCATCGTGGACCTGCTGTTCAA GACCAACCGGAAAGTGACCGTGAAGCAGCTGAAAGAGGACTACTTCAAGAAAAT CGAGTGCTTCGACTCCGTGGAAATCTCCGGCGTGGAAGATCGGTTCAACGCCTCC CTGGGCACATACCACGATCTGCTGAAAATTATCAAGGACAAGGACTTCCTGGACAA TGAGGAAAACGAGGACATTCTGGAAGATATCGTGCTGACCCTGACACTGTTTGAG GACAGAGAGATGATCGAGGAACGGCTGAAAACCTATGCCCACCTGTTCGACGACA AAGTGATGAAGCAGCTGAAGCGGCGGAGATACACCGGCTGGGGCAGGCTGAGCC GGAAGCTGATCAACGGCATCCGGGACAAGCAGTCCGGCAAGACAATCCTGGATTT CCTGAAGTCCGACGGCTTCGCCAACAGAAACTTCATGCAGCTGATCCACGACGAC AGCCTGACCTTTAAAGAGGACATCCAGAAAGCCCAGGTGTCCGGCCAGGGCGATA GCCTGCACGAGCACATTGCCAATCTGGCCGGCAGCCCCGCCATTAAGAAGGGCAT CCTGCAGACAGTGAAGGTGGTGGACGAGCTCGTGAAAGTGATGGGCCGGCACAA GCCCGAGAACATCGTGATCGAAATGGCCAGAGAGAACCAGACCACCCAGAAGGG ACAGAAGAACAGCCGCGAGAGAATGAAGCGGATCGAAGAGGGCATCAAAGAGCT GGGCAGCCAGATCCTGAAAGAACACCCCGTGGAAAACACCCAGCTGCAGAACGA GAAGCTGTACCTGTACTACCTGCAGAATGGGCGGGATATGTACGTGGACCAGGAA CTGGACATCAACCGGCTGTCCGACTACGATGTGGACCATATCGTGCCTCAGAGCTT TCTGAAGGACGACTCCATCGACAACAAGGTGCTGACCAGAAGCGACAAGAACCG GGGCAAGAGCGACAACGTGCCCTCCGAAGAGGTCGTGAAGAAGATGAAGAACTA CTGGCGGCAGCTGCTGAACGCCAAGCTGATTACCCAGAGAAAGTTCGACAATCTG ACCAAGGCCGAGAGAGGCGGCCTGAGCGAACTGGATAAGGCCGGCTTCATCAAG AGACAGCTGGTGGAAACCCGGCAGATCACAAAGCACGTGGCACAGATCCTGGAC TCCCGGATGAACACTAAGTACGACGAGAATGACAAGCTGATCCGGGAAGTGAAAG TGATCACCCTGAAGTCCAAGCTGGTGTCCGATTTCCGGAAGGATTTCCAGTTTTAC AAAGTGCGCGAGATCAACAACTACCACCACGCCCACGACGCCTACCTGAACGCCG TCGTGGGAACCGCCCTGATCAAAAAGTACCCTAAGCTGGAAAGCGAGTTCGTGTA CGGCGACTACAAGGTGTACGACGTGCGGAAGATGATCGCCAAGAGCGAGCAGGA AATCGGCAAGGCTACCGCCAAGTACTTCTTCTACAGCAACATCATGAACTTTTTCAA GACCGAGATTACCCTGGCCAACGGCGAGATCCGGAAGCGGCCTCTGATCGAGAC AAACGGCGAAACCGGGGAGATCGTGTGGGATAAGGGCCGGGATTTTGCCACCGT GCGGAAAGTGCTGAGCATGCCCCAAGTGAATATCGTGAAAAAGACCGAGGTGCAG ACAGGCGGCTTCAGCAAAGAGTCTATCCTGCCCAAGAGGAACAGCGATAAGCTGA TCGCCAGAAAGAAGGACTGGGACCCTAAGAAGTACGGCGGCTTCGACAGCCCCA CCGTGGCCTATTCTGTGCTGGTGGTGGCCAAAGTGGAAAAGGGCAAGTCCAAGAA ACTGAAGAGTGTGAAAGAGCTGCTGGGGATCACCATCATGGAAAGAAGCAGCTTC GAGAAGAATCCCATCGACTTTCTGGAAGCCAAGGGCTACAAAGAAGTGAAAAAGG ACCTGATCATCAAGCTGCCTAAGTACTCCCTGTTCGAGCTGGAAAACGGCCGGAA GAGAATGCTGGCCTCTGCCGGCGAACTGCAGAAGGGAAACGAACTGGCCCTGCC CTCCAAATATGTGAACTTCCTGTACCTGGCCAGCCACTATGAGAAGCTGAAGGGCT CCCCCGAGGATAATGAGCAGAAACAGCTGTTTGTGGAACAGCACAAGCACTACCT GGACGAGATCATCGAGCAGATCAGCGAGTTCTCCAAGAGAGTGATCCTGGCCGAC GCTAATCTGGACAAAGTGCTGTCCGCCTACAACAAGCACCGGGATAAGCCCATCA GAGAGCAGGCCGAGAATATCATCCACCTGTTTACCCTGACCAATCTGGGAGCCCCT GCCGCCTTCAAGTACTTTGACACCACCATCGACCGGAAGAGGTACACCAGCACCA AAGAGGTGCTGGACGCCACCCTGATCCACCAGAGCATCACCGGCCTGTACGAGAC ACGGATCGACCTGTCTCAGCTGGGAGGCGACAAAAGGCCGGCGGCCACGAAAAA GGCCGGCCAGGCAAAAAAGAAAAAGGAATTCGGCAGTGGAGAGGGCAGAGGAA GTCTGCTAACATGCGGTGACGTCGAGGAGAATCCTGGCCCAGGTACCCCTAGGGG ATCCACCGGATCTAGATAACTGATCATAATCAGCCATACCACATTTGTAGAGGTTTTA CTTGCTTTAAAAAACCTCCCACACCTCCCCCTGAACCTGAAACATAAAATGAATGC AATTGTTGTTGTTAACTTGTTTATTGCAGCTTATAATGGTTACAAATAAAGCAATAGC ATCACAAATTTCACAAATAAAGCATTTTTTTCACTGCATTCTAGTTGTGGTTTGTCCA AACTCATCAATGTATCTTA

### >HOsite-TMEP: HO cut sites + Mini-TK promoter + ZsGreen + SV40 poly(A) signal

TTTATGGGACTACTTCGCGCAACAGTATAATTCGCATATTAAGGTGACGCGTGTGGC CTCGAACACCGAGCGACCCTGCAGCGACCCGCTTAAAAGATCCGCCACCATGGCC CAGTCCAAGCACGGCCTGACCAAGGAGATGACCATGAAGTACCGCATGGAGGGC TGCGTGGACGGCCACAAGTTCGTGATCACCGGCGAGGGCATCGGCTACCCCTTCA AGGGCAAGCAGGCCATCAACCTGTGCGTGGTGGAGGGCGGCCCCTTGCCCTTCG CCGAGGACATCTTGTCCGCCGCCTTCATGTACGGCAACCGCGTGTTCACCGAGTA CCCCCAGGACATCGTCGACTACTTCAAGAACTCCTGCCCCGCCGGCTACACCTGG GACCGCTCCTTCCTGTTCGAGGACGGCGCCGTGTGCATCTGCAACGCCGACATCA CCGTGAGCGTGGAGGAGAACTGCATGTACCACGAGTCCAAGTTCTACGGCGTGAA CTTCCCCGCCGACGGCCCCGTGATGAAGAAGATGACCGACAACTGGGAGCCCTC CTGCGAGAAGATCATCCCCGTGCCCAAGCAGGGCATCTTGAAGGGCGACGTGAG CATGTACCTGCTGCTGAAGGACGGTGGCCGCTTGCGCTGCCAGTTCGACACCGTG TACAAGGCCAAGTCCGTGCCCCGCAAGATGCCCGACTGGCACTTCATCCAGCACA AGCTGACCCGCGAGGACCGCAGCGACGCCAAGAACCAGAAGTGGCACCTGACC GAGCACGCCATCGCCTCCGGCTCCGCCTTGCCCTAAGGATCCACCGGATCTAGATA ACTGATCATAATCAGCCATACCACATTTGTAGAGGTTTTACTTGCTTTAAAAAACCTC CCACACCTCCCCCTGAACCTGAAACATAAAATGAATGCAATTGTTGTTGTTAACTTG TTTATTGCAGCTTATAATGGTTACAAATAAAGCAATAGCATCACAAATTTCACAAATA AAGCATTTTTTTCACTGCATTCTAGTTGTGGTTTGTCCAAACTCATCAATGTATCTTA

### >HOsite-TMCP: HO cut sites + Mini-TK promoter + SV40 NLS + Cas9 + SV40 NLS + SV40 poly(A) signal

TTTATGGGACTACTTCGCGCAACAGTATAATTCGCATATTAAGGTGACGCGTGTGGC CTCGAACACCGAGCGACCCTGCAGCGACCCGCTTAAAAGATCCAAGCTTGCCACC ATGGACTATAAGGACCACGACGGAGACTACAAGGATCATGATATTGATTACAAAGA CGATGACGATAAGATGGCCCCAAAGAAGAAGCGGAAGGTCGGTATCCACGGAGTC CCAGCAGCCGACAAGAAGTACAGCATCGGCCTGGACATCGGCACCAACTCTGTGG GCTGGGCCGTGATCACCGACGAGTACAAGGTGCCCAGCAAGAAATTCAAGGTGCT GGGCAACACCGACCGGCACAGCATCAAGAAGAACCTGATCGGAGCCCTGCTGTT CGACAGCGGCGAAACAGCCGAGGCCACCCGGCTGAAGAGAACCGCCAGAAGAA GATACACCAGACGGAAGAACCGGATCTGCTATCTGCAAGAGATCTTCAGCAACGA GATGGCCAAGGTGGACGACAGCTTCTTCCACAGACTGGAAGAGTCCTTCCTGGTG GAAGAGGATAAGAAGCACGAGCGGCACCCCATCTTCGGCAACATCGTGGACGAG GTGGCCTACCACGAGAAGTACCCCACCATCTACCACCTGAGAAAGAAACTGGTGG ACAGCACCGACAAGGCCGACCTGCGGCTGATCTATCTGGCCCTGGCCCACATGAT CAAGTTCCGGGGCCACTTCCTGATCGAGGGCGACCTGAACCCCGACAACAGCGA CGTGGACAAGCTGTTCATCCAGCTGGTGCAGACCTACAACCAGCTGTTCGAGGAA AACCCCATCAACGCCAGCGGCGTGGACGCCAAGGCCATCCTGTCTGCCAGACTG AGCAAGAGCAGACGGCTGGAAAATCTGATCGCCCAGCTGCCCGGCGAGAAGAAG AATGGCCTGTTCGGAAACCTGATTGCCCTGAGCCTGGGCCTGACCCCCAACTTCA AGAGCAACTTCGACCTGGCCGAGGATGCCAAACTGCAGCTGAGCAAGGACACCT ACGACGACGACCTGGACAACCTGCTGGCCCAGATCGGCGACCAGTACGCCGACC TGTTTCTGGCCGCCAAGAACCTGTCCGACGCCATCCTGCTGAGCGACATCCTGAG AGTGAACACCGAGATCACCAAGGCCCCCCTGAGCGCCTCTATGATCAAGAGATAC GACGAGCACCACCAGGACCTGACCCTGCTGAAAGCTCTCGTGCGGCAGCAGCTG CCTGAGAAGTACAAAGAGATTTTCTTCGACCAGAGCAAGAACGGCTACGCCGGCT ACATTGACGGCGGAGCCAGCCAGGAAGAGTTCTACAAGTTCATCAAGCCCATCCT GGAAAAGATGGACGGCACCGAGGAACTGCTCGTGAAGCTGAACAGAGAGGACCT GCTGCGGAAGCAGCGGACCTTCGACAACGGCAGCATCCCCCACCAGATCCACCT GGGAGAGCTGCACGCCATTCTGCGGCGGCAGGAAGATTTTTACCCATTCCTGAAG GACAACCGGGAAAAGATCGAGAAGATCCTGACCTTCCGCATCCCCTACTACGTGG GCCCTCTGGCCAGGGGAAACAGCAGATTCGCCTGGATGACCAGAAAGAGCGAGG AAACCATCACCCCCTGGAACTTCGAGGAAGTGGTGGACAAGGGCGCTTCCGCCC AGAGCTTCATCGAGCGGATGACCAACTTCGATAAGAACCTGCCCAACGAGAAGGT GCTGCCCAAGCACAGCCTGCTGTACGAGTACTTCACCGTGTATAACGAGCTGACC AAAGTGAAATACGTGACCGAGGGAATGAGAAAGCCCGCCTTCCTGAGCGGCGAG CAGAAAAAGGCCATCGTGGACCTGCTGTTCAAGACCAACCGGAAAGTGACCGTG AAGCAGCTGAAAGAGGACTACTTCAAGAAAATCGAGTGCTTCGACTCCGTGGAAA TCTCCGGCGTGGAAGATCGGTTCAACGCCTCCCTGGGCACATACCACGATCTGCT GAAAATTATCAAGGACAAGGACTTCCTGGACAATGAGGAAAACGAGGACATTCTG GAAGATATCGTGCTGACCCTGACACTGTTTGAGGACAGAGAGATGATCGAGGAAC GGCTGAAAACCTATGCCCACCTGTTCGACGACAAAGTGATGAAGCAGCTGAAGCG GCGGAGATACACCGGCTGGGGCAGGCTGAGCCGGAAGCTGATCAACGGCATCCG GGACAAGCAGTCCGGCAAGACAATCCTGGATTTCCTGAAGTCCGACGGCTTCGCC AACAGAAACTTCATGCAGCTGATCCACGACGACAGCCTGACCTTTAAAGAGGACA TCCAGAAAGCCCAGGTGTCCGGCCAGGGCGATAGCCTGCACGAGCACATTGCCAA TCTGGCCGGCAGCCCCGCCATTAAGAAGGGCATCCTGCAGACAGTGAAGGTGGTG GACGAGCTCGTGAAAGTGATGGGCCGGCACAAGCCCGAGAACATCGTGATCGAA ATGGCCAGAGAGAACCAGACCACCCAGAAGGGACAGAAGAACAGCCGCGAGAG AATGAAGCGGATCGAAGAGGGCATCAAAGAGCTGGGCAGCCAGATCCTGAAAGA ACACCCCGTGGAAAACACCCAGCTGCAGAACGAGAAGCTGTACCTGTACTACCTG CAGAATGGGCGGGATATGTACGTGGACCAGGAACTGGACATCAACCGGCTGTCCG ACTACGATGTGGACCATATCGTGCCTCAGAGCTTTCTGAAGGACGACTCCATCGAC AACAAGGTGCTGACCAGAAGCGACAAGAACCGGGGCAAGAGCGACAACGTGCC CTCCGAAGAGGTCGTGAAGAAGATGAAGAACTACTGGCGGCAGCTGCTGAACGC CAAGCTGATTACCCAGAGAAAGTTCGACAATCTGACCAAGGCCGAGAGAGGCGG CCTGAGCGAACTGGATAAGGCCGGCTTCATCAAGAGACAGCTGGTGGAAACCCG GCAGATCACAAAGCACGTGGCACAGATCCTGGACTCCCGGATGAACACTAAGTAC GACGAGAATGACAAGCTGATCCGGGAAGTGAAAGTGATCACCCTGAAGTCCAAGC TGGTGTCCGATTTCCGGAAGGATTTCCAGTTTTACAAAGTGCGCGAGATCAACAAC TACCACCACGCCCACGACGCCTACCTGAACGCCGTCGTGGGAACCGCCCTGATCA AAAAGTACCCTAAGCTGGAAAGCGAGTTCGTGTACGGCGACTACAAGGTGTACGA CGTGCGGAAGATGATCGCCAAGAGCGAGCAGGAAATCGGCAAGGCTACCGCCAA GTACTTCTTCTACAGCAACATCATGAACTTTTTCAAGACCGAGATTACCCTGGCCAA CGGCGAGATCCGGAAGCGGCCTCTGATCGAGACAAACGGCGAAACCGGGGAGAT CGTGTGGGATAAGGGCCGGGATTTTGCCACCGTGCGGAAAGTGCTGAGCATGCCC CAAGTGAATATCGTGAAAAAGACCGAGGTGCAGACAGGCGGCTTCAGCAAAGAGT CTATCCTGCCCAAGAGGAACAGCGATAAGCTGATCGCCAGAAAGAAGGACTGGGA CCCTAAGAAGTACGGCGGCTTCGACAGCCCCACCGTGGCCTATTCTGTGCTGGTG GTGGCCAAAGTGGAAAAGGGCAAGTCCAAGAAACTGAAGAGTGTGAAAGAGCTG CTGGGGATCACCATCATGGAAAGAAGCAGCTTCGAGAAGAATCCCATCGACTTTCT GGAAGCCAAGGGCTACAAAGAAGTGAAAAAGGACCTGATCATCAAGCTGCCTAAG TACTCCCTGTTCGAGCTGGAAAACGGCCGGAAGAGAATGCTGGCCTCTGCCGGCG AACTGCAGAAGGGAAACGAACTGGCCCTGCCCTCCAAATATGTGAACTTCCTGTA CCTGGCCAGCCACTATGAGAAGCTGAAGGGCTCCCCCGAGGATAATGAGCAGAAA CAGCTGTTTGTGGAACAGCACAAGCACTACCTGGACGAGATCATCGAGCAGATCA GCGAGTTCTCCAAGAGAGTGATCCTGGCCGACGCTAATCTGGACAAAGTGCTGTC CGCCTACAACAAGCACCGGGATAAGCCCATCAGAGAGCAGGCCGAGAATATCATC CACCTGTTTACCCTGACCAATCTGGGAGCCCCTGCCGCCTTCAAGTACTTTGACAC CACCATCGACCGGAAGAGGTACACCAGCACCAAAGAGGTGCTGGACGCCACCCT GATCCACCAGAGCATCACCGGCCTGTACGAGACACGGATCGACCTGTCTCAGCTG GGAGGCGACAAAAGGCCGGCGGCCACGAAAAAGGCCGGCCAGGCAAAAAAGAA AAAGGAATTCGGCAGTGGAGAGGGCAGAGGAAGTCTGCTAACATGCGGTGACGT CGAGGAGAATCCTGGCCCAGGTACCCCTAGGGGATCCACCGGATCTAGATAACTG ATCATAATCAGCCATACCACATTTGTAGAGGTTTTACTTGCTTTAAAAAACCTCCCAC ACCTCCCCCTGAACCTGAAACATAAAATGAATGCAATTGTTGTTGTTAACTTGTTTAT TGCAGCTTATAATGGTTACAAATAAAGCAATAGCATCACAAATTTCACAAATAAAGC ATTTTTTTCACTGCATTCTAGTTGTGGTTTGTCCAAACTCATCAATGTATCTTA

